# Sequencing, de novo assembly of Ludwigia plastomes, and comparative analysis within the Onagraceae family

**DOI:** 10.1101/2023.10.20.563230

**Authors:** F. Barloy-Hubler, A.-L. Le Gac, C. Boury, E. Guichoux, D. Barloy

## Abstract

The Onagraceae family, which belongs to the order Myrtales, consists of approximately 657 species and 17 genera, including the genus *Ludwigia* L., which is comprised of 82 species. There are few genomic resources for Onagraceae, which limits phylogenetic and population genetics, as well as genomic studies. In this study, new complete plastid genomes of Ludwigia grandiflora subps. hexapetala (Lgh) and Ludwigia peploides subsp montevidensis (Lpm) were generated using a combination of different sequencing technologies. These plastomes were then compared to the published Ludwigia octovalvis (Lo) plastid genome, which was re-annotated by the authors. We initially sequenced and assembled the chloroplast (cp) genomes of Lpm and Lgh using a hybrid strategy. We observed the existence of two Lgh haplotypes and two potential Lpm haplotypes. Lgh, Lpm, and Lo plastomes were similar in terms of genome size, gene number, structure, and inverted repeat (IR) boundaries, comparable to other species in the Myrtales order. A total of 45 to 65 SSRs (*simple sequence* repeats), were detected, depending on the species, with the majority consisting solely of A and T, which is common among angiosperms. Four chloroplast genes (matK, accD, ycf2 and ccsA) were found under positive selection pressure, which is commonly associated with plant development, and especially in aquatic plants such as Lgh, and Lpm. Our hybrid sequencing approach revealed the presence of two Lgh plastome haplotypes which will help to advance phylogenetic and evolutionary studies, not only specifically for Ludwigia, but also the Onagraceae family and Myrtales order. To enhance the robustness of our findings, a larger dataset of chloroplast genomes would be beneficial.

## Introduction

The Onagraceae family belongs to the order Myrtales which includes approximately 657 species of herbs, shrubs, and trees across 17 genera grouped into two subfamilies: subfam. Ludwigioideae W. L. Wagner and Hoch, which only has one genus (Ludwigia L.), and subfam. Onagroideae which contains six tribes and 21 genera (Wagner, Hoch, and Raven 2007a). Ludwigia L. is composed of 83 species(Levin et al. 2003)(Levin et al. 2004). The current classification for Ludwigia L., which are composed of several hybrid and/or polyploid species, lists 23 sections. A recent molecular analysis is clarified and supported several major relationships in the genus but has challenged the complex sectional classification of Ludwigia L.(S. H. Liu et al. 2017).

The diploid species Ludwigia peploides (Kunth) Raven subsp. montevidensis (Spreng.) (Raven P.H. 1963) (named here Lpm) (2n=16), and the decaploid species, Ludwigia grandiflora (Michx) Greuter & Burdet subsp. hexapetala (Hook. & Arn) Nesom & Kartesz (named here Lgh) (2n=80), reproduce essentially by clonal propagation, which suggests that there is a low genetic diversity within the species (Dandelot et al. 2005). Lgh and Lpm are native to South America and are considered as one of the most aggressive aquatic invasive plants (Reddy et al., n.d.). Largely distributed in aquatic environments in North America and in Europe (Hussner, Windhaus, and Starfinger 2016), both species have been found to degrade major watersheds as well as aquatic and riparian ecosystems (Grewell, Netherland, and Skaer Thomason 2016) leading ecological and economical damages. In France, both species occupied aquatic habitats, such as static or slow-flowing waters, riversides, and have recently been observed in wet meadows (Lambert et al. 2010). The transition from an aquatic to a terrestrial habitat has led to the emergence of two Lgh morphotypes (Haury et al. 2014a). The appearance of metabolic and morphological adaptations could explain the ability to acclimatize to terrestrial conditions, and this phenotypic plasticity involves various genomic and epigenetic modifications (Kevin Billet et al. 2018).

Adequate genomic resources are necessary in order to be identify the genes and metabolic pathways involved in the adaptation process leading to plant invasion (Gioria et al. 2023) with genomic information making it possible to predict and control invasiveness (Moravcová et al. 2015). However, even though the number of terrestrial plant genomes has increased considerably over the last 20 years, only a small fraction (∼ 0.16%) have been sequenced, with some clades being significantly more represented than others (Marks et al. 2021). Thus, for the Onagraceae family (which includes Ludwigia sp.), only a handful of chloroplast sequences (plastomes) are available, and the complete genome has not yet been sequenced. If Lpm is a diploid species (2n=2x=16) with a relatively small genome size (262 Mb), Lgh is a decaploid species (2n=10x=80) with a large size genome of 1419 Mb (Barloy et al. 2024). Obtaining a reference genome for these two non-model species without having a genome close to the Ludwigia species is challenging and development of plastome and/or mitogenome will be a first step to generate genomic resource. As of April 2023, there are 10,712 reference plastomes listed on GenBank (Release 255: April 15 2023), with the vast majority (10,392 genomes) belonging to Viridiplantae (green plants). However, in release 255, the number of plastomes available for the Onagraceae family is limited, with only 36 plastomes currently listed. Among these, 15 plastomes are from the tribe Epilobieae, with 11 in the Epilobium genus and 4 in the Chamaenerion genus. Additionally, there are 23 plastomes from the tribe Onagreae, with 17 in the Oenothera genus, 5 in the Circaea genus, and only one in the Ludwigia genus. The Ludwigia octovalvis chloroplast genome was released in 2016 as a unique haplotype of approximately 159 kb (S. H. Liu et al. 2016). L. octovalvis belongs to sect. Macrocarpon (Micheli) H.Hara while Lpm and Lgh belong to Jussieae section (Zardini and Raven 1992)(Hoch, Wagner, and Raven 2015). Generally, the inheritance of chloroplast genomes is considered to be maternal in angiosperms. However, biparentally inherited chloroplast genomes could potentially exist in approximately 20% of angiosperm species (Hu et al. 2008)(Q. Zhang and Sodmergen 2010). Both maternal and biparental inheritance are described in the Onagraceae family. In tribe Onagreae, Oenothera subsect. Oenothera are known to have biparental plastid inheritance (Wagner, Hoch, and Raven 2007b)(Jones and Cleland 1974). In tribe Epilobieae, biparental plastid inheritance was also reported in Epilobium L. with mainly maternal transmission, and very low proportions of paternally transmitted chloroplasts (Schmitz and Kowallik 1986).

The chloroplast is the symbolic organelle of plants and plays a fundamental role in photosynthesis. Chloroplasts evolved from cyanobacteria through endosymbiosis and thereby inherited components of photosynthesis reactions (photosystems, electron transfer and ATP synthase) and gene expression systems (transcription and translation)(Sato 2021). In general, chloroplast genomes (plastomes) are highly conserved in size, structure, and genetic content. They are rather small (120-170 kb,(Gualberto et al. 2014)), with a quadripartite structure comprising two long identical inverted repeats (IR, 10–30 kb) separated by large and a small single copy regions (LSC and SSC, respectively). They are also rich in genes, with around 100 unique genes encoding key proteins involved in photosynthesis, and a comprehensive set of ribosomal RNAs (rRNAs) and transfer RNAs (tRNAs)(Tonti-Filippini et al. 2017). Plastomes are generally circular but linear shapes also exist (Oldenburg and Bendich 2016). Chloroplast DNA usually represents 5-20% of total DNA extracted from young leaves and therefore low-coverage whole genome sequencing can generate enough data to assemble an entire chloroplast genome (Twyford and Ness 2017).

If we refer to their GenBank records, more than 95% of these plastomes were sequenced by so-called short read techniques (mostly Illumina). However, in most seed plants, the plastid genome exhibits two large inverted repeat regions (60 to 335 kb,(Twyford and Ness 2017)), which are longer than the short read lengths (< 300 bp). This leads to incomplete or approximate assemblies (Wang et al. 2018a). Recent long-read sequencing (> 1000 bp) provides compelling evidence that terrestrial plant plastomes exhibit two structural haplotypes. These haplotypes are present in equal proportions and differ in their inverted repeat (IR) orientation (Wang, Lanfear, and Gaut 2019). This shows the importance of using the so-called third generation sequence (TGS, PacBio or Nanopore) to correctly assemble the IRs of chloroplasts and to identify any different structural haplotypes. The current problem with PacBio or Nanopore long read sequencing is the higher error rate compared to short read technology (Ferrarini et al. 2013)(Jain et al. 2018)(Rang, Kloosterman, and de Ridder 2018). Thus, a hybrid strategy which combines long reads (to access the genomic structure) and short reads (to correct sequencing errors) could be effective (Wang et al. 2018a)(Scheunert et al. 2020).

Here, we report the newly sequenced complete plastid genomes of Ludwigia grandiflora subps. hexapetala (Lgh) and Ludwigia peploides subsp montevidensis (Lpm), using a combination of different sequencing technologies, as well as a re-annotated comparative genomic analysis of the published Ludwigia octovalvis (Lo) plastid. The main objectives of this study are (1) to assemble and annotate the plastomes of two new species of Ludwigia sp., (2) to reveal the divergent sequence hotspots of the plastomes in this genus and in the Onagraceae (3) to identify the genes under positive selection.

To achieve this, we utilized long read sequencing data from Oxford Nanopore and short read sequencing data from Illumina to assemble the Lgh plastomes and compared these assemblies with those obtained solely from long reads of Lpm. We also compared both plastomes to the published plastome of Lo. Our findings demonstrated the value of de novo assembly in reducing assembly errors and enabling accurate reconstruction of full heteroplasmy. We also evaluated the performance of a variety of software for sequence assembly and correction in order to define a workflow that will be used in the future to assemble Ludwigia sp. mitochlondrial and nuclear genomes. Finally, the three new Ludwigia plastomes generated by our study make it possible to extend the phylogenetic study of the Onagraceae family and to compare it with previously published analyses (S. H. Liu et al. 2017)(Bedoya and Madriñán 2015)(S. H. Liu et al. 2020).

## Material and Methods

### Plant sampling and experimental design

The original plant materials were collected in June of 2018 near to Nantes (France) and formal identified by D. Barloy. *L. grandiflora* subsp*. hexapetala (Lgh)* plants were taken from the Mazerolles swamps (N47 23.260, W1 28.206), and *L. peploides* subsp*. montevidensis* (*Lpm*) plants from La Musse (N 47.240926, W −1.788688)). Plants were cultivated in a growth chamber in a mixture of ^1^/_3_ soil, ^1^/_3_ sand, ^1^/_3_ loam with flush water level, at 22°C and a 16 h/8 h (light/dark) cycle. A single stem of 10 cm for each species was used for vegetative propagation in order to avoid potential genetic diversity. *De novo* shoots, taken three centimeters from the apex, were sampled for each species. Samples for gDNA extraction were pooled and immediately snap-frozen in liquid nitrogen, then lyophilized over 48 h using a Cosmos 20K freeze-dryer (Cryotec, Saint-Gély-du-Fesc, France) and stored at room temperature. All the plants were destroyed after being used as required by French authorities for invasive plants (article 3, prefectorial decree n°2018/SEE/2423).

Due to high polysaccharide content and polyphenols in *Lpm* and *Lgh* tissues and as no standard kit provided good DNA quality for sequencing, genomic DNA extraction was carried out using a modified version of the protocol proposed by Panova et al in 2016, with three purification steps (Panova et al. 2016).

40 mg of lyophilized buds were ground at 30 Hz for 60 s (Retsch MM200 mixer mill, FISHER). The ground tissues were lysed with 1 ml CF lysis buffer (MACHEREY-NAGEL) supplemented with <u>20 μl</u> RNase and incubated for 1 h at 65°C under agitation. 20 <u>μ</u>l proteinase K was then added before another incubation for 1 h at 65°C under agitation. To avoid breaking the DNA during pipetting, the extracted DNA was recovered using a Phase-lock gel tube as described in Belser (Belser et al. 2018). The extracts were transferred to 2 ml tubes containing phase-lock gel, and an equal volume of PCIA (Phenol, Chloroform, Isoamyl Alcohol; 25:24:1) was added. After shaking for 5 min, tubes were centrifuged at 11000 g for 20 min. The aqueous phase was transferred into a new tube containing phase-lock gel and extraction with PCIA was repeated. DNA was then precipitated after addition of an equal volume of binding buffer C4 (MACHEREY-NAGEL) and 99% ethanol overnight at 4°C or 1 h in ice then centrifuged at 800 rpm for 10 min. After removal of the supernatant, 1 ml of CQW buffer was added then the pellet of DNA was re-suspended. Next, DNA purification was carried out by adding a 2 ml mixture of wash buffer PW2 (MACHEREY-NAGEL), wash buffer B5 (MACHEREY-NAGEL), and ethanol at 99% in equal volumes, followed by centrifugation at 800 rpm for 10 min. This DNA purification step was carried out twice. Finally, the DNA pellet was dried in the oven at 70°C for 30 min then re-suspended in 100 <u>μ</u>l elution buffer BE (MACHEREY-NAGEL) (5 mM Tris solution, pH 8.5) after 10 min incubation at 65°C under agitation.

A second purification step was performed using a PCR product extraction from gel agarose kit from Macherey-Nagel (MN) NucleoSpin® Gel and PCR Clean-up kit and restarting the above protocol from the step with the addition of CQW buffer then PW2 buffer.

The third purification step consisted of DNA purification using a Macherey-Nagel (MN) NucleoMag kit for clean-up and size selection. Finally, the DNA was resuspended after a 5 min incubation at 65°C in 5 mM TRIS at pH 8.5.

The quantity and quality of the gDNA was verified using a NanoDrop spectrometer, electrophoresis on agarose gel and ethidium bromide staining under UV light and Fragment Analyzer (Agilent Technologies) of the University of Rennes1.

### Library preparation and sequencing

MiSeq (Illumina) and GridION (Oxford Nanopore Technologies, referred to here as ONT) sequencing were performed at the PGTB (doi:10.15454/1.5572396583599417E12). *Lgh* and *Lpm* genomic DNA were re-purified using homemade SPRI beads (1.8X ratio)*. Lgh* has a large genome size of 1419 Mb, 5-fold larger than *Lpm* genome 262 Mb (Barloy et al. 2024). SR (Illumina, one run) and LR (Oxford Nanopore, three runs) sequencing were therefore carried out for *Lgh* and only LR sequencing for *Lpm* (one run). For Illumina sequencing, 200 ng of *Lgh* DNA was used according to the QIAseq FX DNA Library Kit protocol (Qiagen). The final library was checked on TapeStation D5000 screentape (Agilent Technologies) and quantified using a QIAseq Library Quant Assay Kit (Qiagen). The pool was sequenced on an Illumina MiSeq using V3 chemistry and 600 cycles (2×300bp). For ONT sequencing, around 8 µg of *Lgh* and *Lpm* DNA were size selected using a Circulomics SRE kit (according to the manufacturer’s instructions) before library preparation using a SQK-LSK109 ligation sequencing kit following ONT recommendations. Basecalling in High Accuracy - Guppy version: 4.0.11 (MinKNOW GridION release 20.06.9) was performed for the 48 h of sequencing. Long reads (LR) and short reads (SR) were available for *Lgh* and only LR for *Lpm*.

#### Chloroplast assemblies

Quality controls and preprocessing of sequences were conducted using Guppy v4.0.14 for long reads (via Oxford Nanopore Technology Client access) and fastp v0.20.0 (S. Chen et al. 2018) for short reads, using Q15, since increasing the Phred quality to 20 or higher has no effect on the number of sequences retained (66%). A preliminary draft assembly was performed using *Lgh* short-reads (SR, 2*23,067,490 reads) with GetOrganelle v1.7.0 (Jin et al. 2020) and NOVOPlasty v4.2.1(Dierckxsens, Mardulyn, and Smits 2017), and chloroplastic short and long reads were extracted by mapping against this draft genome. Chloroplastic short reads were then *de novo* assemble using Velvet (version 1.2.10) (Zerbino and Birney 2008), ABySS (version 2.1.5 (Simpson et al. 2009)(Jackman et al. 2017)), MEGAHIT (1.1.2,(D. Li et al. 2016)), and SPAdes (version 3.15.4,(Bankevich et al. 2012)), without and with prior error correction. The best k-mer parameters were tested using kmergenie (Chikhi and Medvedev 2014) and k=99 was found to be optimal. For ONT reads, *Lgh* (550,516 reads) and *Lpm* (68,907 reads) reads were self-corrected using CANU 1.8 (Koren et al. 2017)or SR-corrected using Ratatosk (Holley et al. 2021) and *de novo* assembly using CANU (Koren et al. 2017) and FLYE 2.8.2 (Kolmogorov et al. 2019) run with the option -- meta and –plasmids. For all these assemblers, unless otherwise specified, we used the default parameters.

#### Post plastome assembly validation

As we used many assemblers and different strategies, we produced multiple contigs that needed to be analyzed and filtered in order to retain only the most robust plastomes. For that, all assemblies were evaluated using the QUality ASsessment Tool (QUAST) for quality assessment (Gurevich et al. 2013) and visualized using BANDAGE (Wick et al. 2015), both using default parameters. BANDAGE compatible graphs (.gfa format) were created with the megahit_toolkit for MEGAHIT (D. Li et al. 2016) and with gfatools for ABySS (Jackman et al. 2017). Overlaps between fragments were manually checked and ambiguous “IUPAC or N” nucleotides were also biocured with Illumina reads when available.

### Chloroplast genome annotation

Plastomes were annotated via the GeSeq (Tillich et al. 2017) using ARAGORN and tRNAscan_SE to predict tRNAs and rRNAs and tRNAscan_SE to predict tRNAs and rRNAs and via Chloe prediction site (X. Zhong 2020). The previously reported *Lo* chloroplast genome was also similarly re-annotated to facilitate genomic comparisons. Gene boundaries, alternative splice isoforms, pseudogenes and gene names and functions were manually checked and biocurated using Geneious (v.10). Finally, plastomes were represented using OrganellarGenomeDRAW (OGDRAW)(Greiner, Lehwark, and Bock 2019). These genomes were submitted to GenBank at the National Center of Biotechnology Information (NCBI) with specific accession numbers (for *Lgh* haplotype 1, (LGH1) OR166254 and *Lgh* haplotype 2, (LGH2) OR166255; for *Lpm* haplotype, (LPM) OR166256) using annotation tables generated through GB2sequin (Lehwark and Greiner 2019).

#### SSRs and Repeat Sequences Analysis

Simple Sequence Repeats (SSRs) were analyzed through the MISA web server (Beier et al. 2017), with parameters set to 10, 5, 4, 3, 3, and 3 for mono-, di-, tri-, tetra-, penta-, and hexa-nucleotides, respectively. Direct, reverse and palindromic repeats were identified using RepEx (Gurusaran, Ravella, and Sekar 2013). Parameters used were: for inverted repeats (min size 15 nt, spacer = local, class = exact); for palindromes (min size 20 nt); for direct repeats (minimum size 30 nt, minimum repeat similarity 97%). Tandem repeats were identified using Tandem Repeats Finder(Benson 1999), with parameters set to two for the alignment parameter match and seven for mismatches and indels. The IRa region was removed for all these analyses to avoid over representation of the repeats.

#### Comparative chloroplast genomic analyses

*Lgh* and *Lpm* plastomes were compared with the reannotated and biocurated *Lo* plastome using mVISTA program (Frazer et al. 2004), with the LAGAN alignment algorithm (Brudno et al. 2003) and a cut-off of 70% identity. Nucleotide diversity (Pi) was analyzed using the software DnaSP v.6.12.01 (J. Rozas and Rozas 1999) (Julio Rozas et al. 2017)with step size set to 200 bp and window length to 300 bp. IRscope (Amiryousefi, Hyvönen, and Poczai 2018) was used for the analyses of inverted repeat (IR) region contraction and expansion at the junctions of chloroplast genomes. To assess the impact of environmental pressures on the evolution of these three *Ludwigia* species, we calculated the nonsynonymous (Ka) and synonymous (Ks) substitutions and their ratios (ω = Ks/Ks) using TBtools (C. Chen et al. 2020) to measure the selective pressure. Genes with ω < 1, ω = 1, and 1 < ω were considered to be under purifying selection (negative selection), neutral selection, and positive selection, respectively.

### Phylogenetic analysis of *Ludwigia* based on MatK sequences

We performed a phylogenetic analysis on the *Ludwigia* genus using the MatK, only protein coding barcode available for a large number of *Ludwigia* species. All MatK amino acid sequences were aligned with the FFT-NS-2 (Fast Fourier Transform-based Narrow Search) algorithm and BLOSUM62 scoring matrix using MAFFT 7 (Katoh et al. 2002). The phylogenetic tree analysis was conducted using the rapid hill-climbing algorithm (command line: -f d) in RAxML 8.2.11 (Stamatakis 2014), with GAMMA JTT (Jones-Taylor-Thornton) protein model. Node support was assessed through fast bootstrapping (-f a) with 1,000 non-parametric bootstrap pseudo-replicates. *Circaea* MatK were selected as outgroup, and all accession numbers are indicated on the phylogenetic tree labels.

### Graphic representation

Statistical analyses were performed using R software in RStudio integrated development environment (R Core Team, 2015, RStudio: Integrated Development for R. RStudio, Inc., Boston, MA, http://www.rstudio.com/). Figures were realized using ggplot2, ggpubr, tidyverse, dplyr, gridExtra, reshape2, and viridis packages. SNPs were represented using trackViewer (Ou and Zhu 2019) and genes represented using gggenes packages.

## Results

### Plastome short read assembly

The chloroplastic fraction of *Lgh* short reads (SR) was extracted by mapping against the two draft haplotypes generated by GetOrganelle, which differ only by a “flip-flop” of the SSC region (Figure 1). Since the assembly by NOVOplasty did not provide any additional information compared to GetOrganelle, it was not retained. This subset (1,360,507 reads) was assembled using ABySS, Velvet, MEGAHIT and SPAdes in order to identify the best assembler for this plant model.

**Figure 1:**
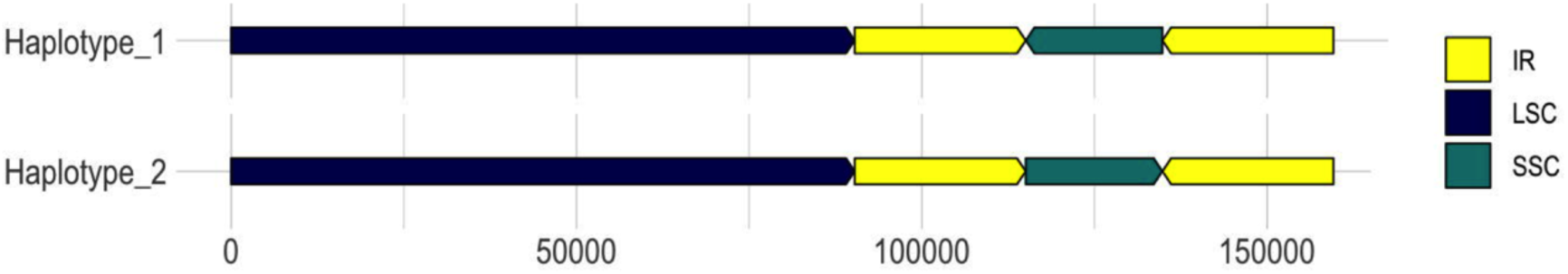
Two structural haplotypes *of L. grandiflora* plastomes representing the flip-flop organization of SSC segment

As shown in Figure 2, both the number and size of contigs depend greatly on the algorithms used and the correction step. The effect of prior read correction is notable for MEGAHIT and Velvet, especially concerning the increase in the size of the large alignment (Add. Figure 1A), loss of misassemblies, and reduction of the number of mismatches (Add. Figure 1B). Investigating results via BANDAGE (Add. Figure 2), we observed that ABySS and SPAdes suggest the tripartite structure with the long single-copy (LSC) region as the larger circle in the graph (blue), joined to the small single-copy region (green) by one copy of the inverted repeats (IRs, red), both IRs being collapsed in a segment of approximately twice the coverage. For Velvet and MEGAHIT, graphs confirm the significant fragmentation of the assemblies, which is improved by prior correction of the reads.

**Figure 2:**
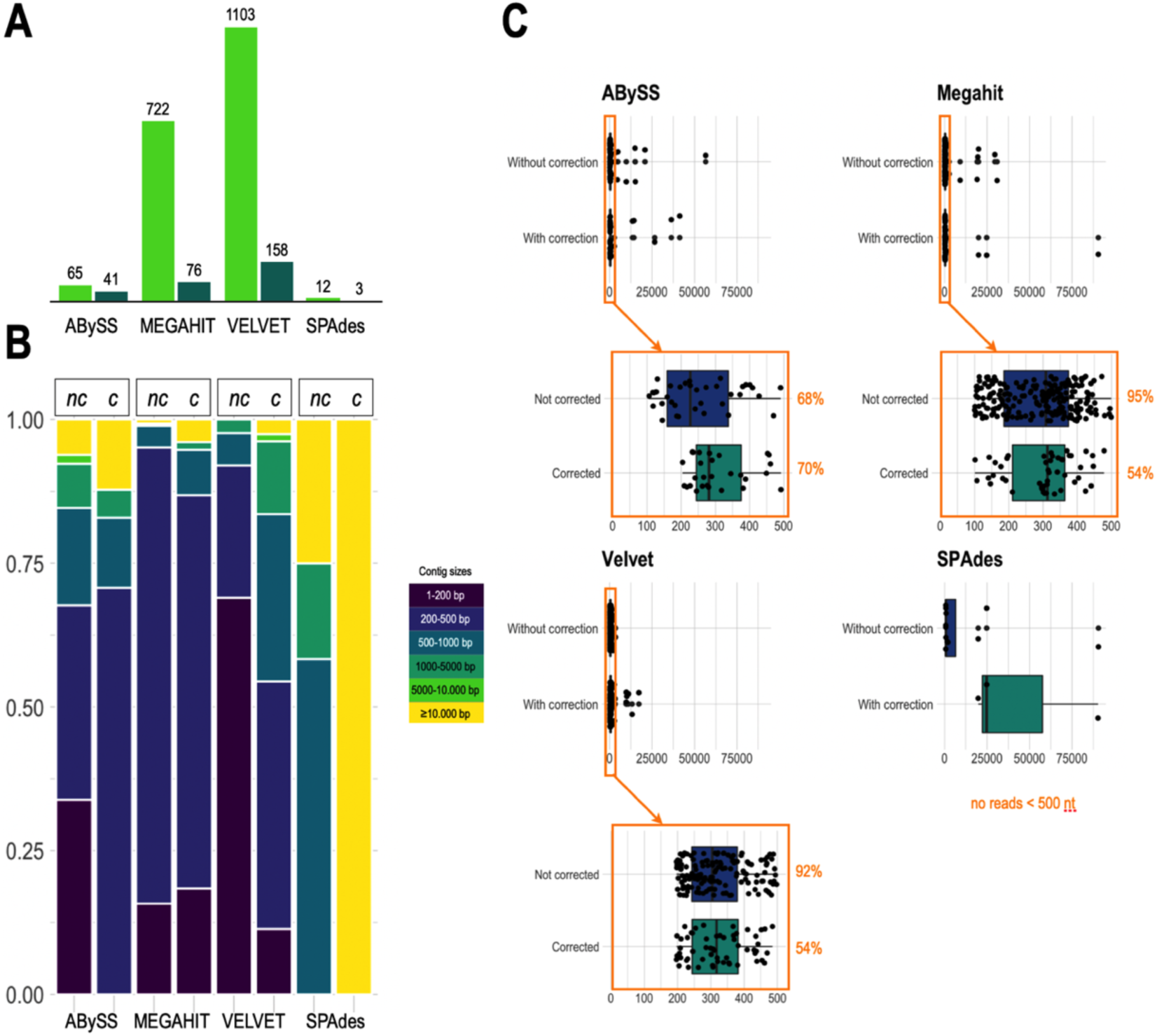
Comparative results of *L. grandiflora* short read (SR) assemblies. **A:** Total number of contigs obtained with the uncorrected (dark green) and corrected (light green) chloroplast SRs for the 4 assemblers (ABySS, MEGAHIT, Velvet and SPAdes). **B:** Comparison of the size of contigs assembled by the 4 tools using corrected or uncorrected SRs. **C:** Boxplot showing the distribution of these contigs by size and the improvement brought by the prior correction of the SRs with the long reads for each tool.

In conclusion, none of the short-read assemblers tested in our study produced a complete plastome. The best result was achieved by SPAdes using corrected short reads (mean coverage 1900 X) to assemble a plastome consisting of three contigs: 90,272 bp (corresponding to LSC), 19,788 bp (corresponding to SSC), and 24,762 bp (corresponding to one of the two copies of the IR).

### Plastome long read assembly

Chloroplast fractions of *Lgh* long reads (28,882 reads) were assembled using CANU or FLYE. With raw data, CANU generates a unique contig (NGA50 112648) corresponding to haplotype 2, whereas FLYE makes two contigs (NGA50 133687) that reconstruct haplotype 1. Self-corrected LR leads to fragmentation into two (CANU) or three (FLYE) contigs which both reconstruct haplotype 1, with a large gap corresponding to one of the IR copies for CANU. Finally, SR-correction by RATATOSK allows CANU to assemble two redundant contigs reproducing haplotype 2 while FLYE makes two contigs corresponding to haplotype 1 (Add. Figure 3A). In conclusion, the two *Lgh* haplotypes were reconstructed (average coverage 700X) and the most complete and accurate hybrid assemblies (99.94% accuracy, Additional Figure 3B) were submitted to GenBank. Unfortunately, due to the absence of short read data, we could only perform self-corrected long read assembly for *Lpm* using CANU. We also compared CANU and FLYE assembler efficiency, and found that assembly using CANU produces 13 contigs whereas FLYE produces 12 contigs. In both cases, only three contigs are required to reconstitute a complete cpDNA assembly (no gap, no N), with an SSC region oriented like those of the *Lgh* haplotype 2 and the *Lo* plastome. Although it is more than likely that these two SSC region orientations also exist for *Lpm*, the low number of nanopore sequences generated (68907 reads) and absence of Illumina short reads prevented us from demonstrating the existence of both haplotypes. As a result, only the “haplotype 2” generated sequence was deposited to Genbank.

**Figure 3:**
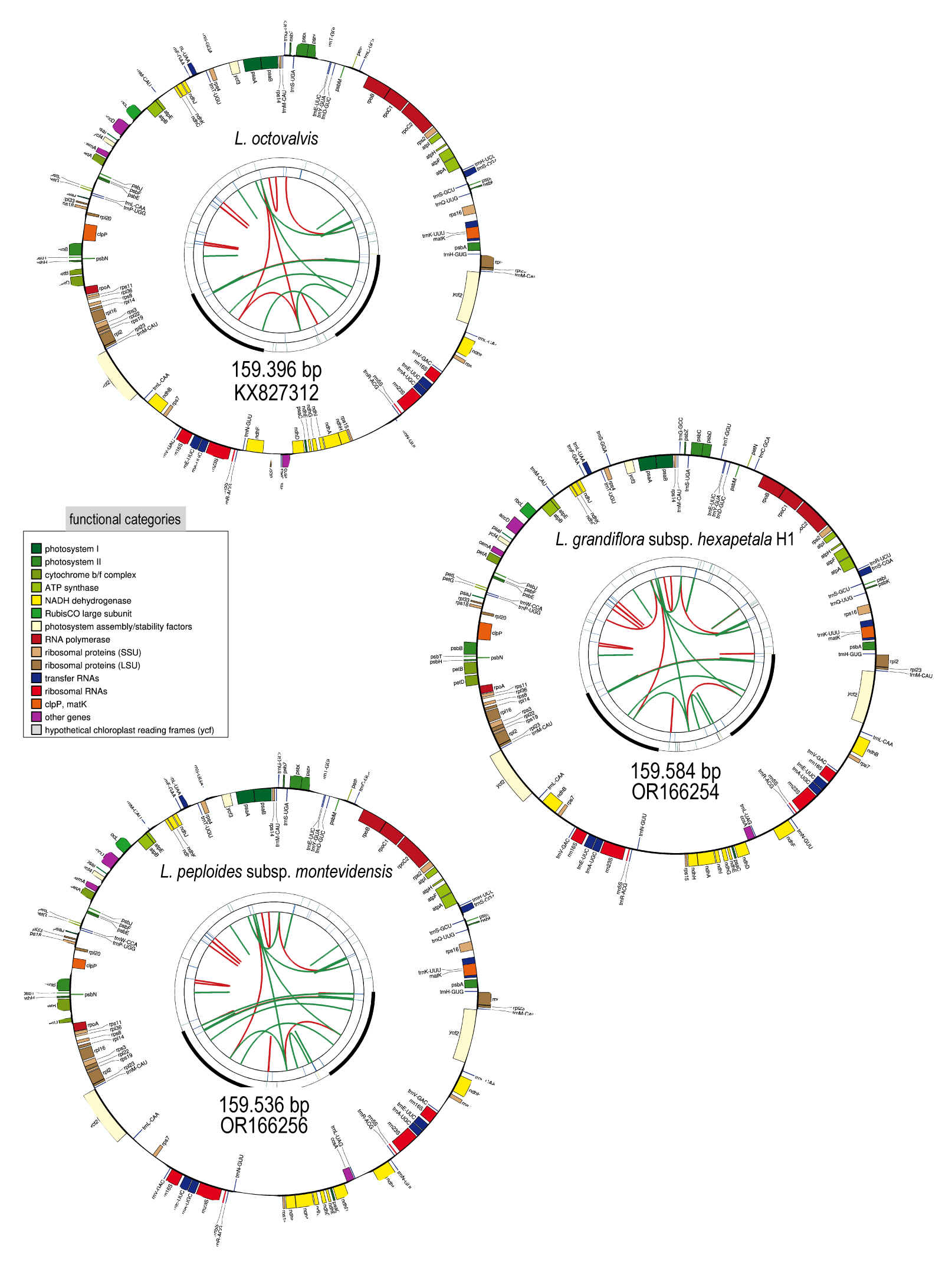
Circular representation of annotations plastomes in *Ludwigia octovalis*, *Ludwigia grandiflora* and *Ludwigia peploides* using ogdraw. Each card contains four circles. From the center outwards, the first circle shows forward and reverse repeats (red and green arcs, respectively). The next circle shows tandem repeats as bars. The third circle shows the microsatellite sequences. Finally, the fourth and fifth circles show the genes colored according to their functional categories (see colored legend). Only the haplotype 1 of *L. grandiflora* is represented as haplotype 2 only diverge by the orientation of the SSC segment. Accession numbers are indicated for each plastome.

### Annotation and comparison of *Ludwigia* plastomes

#### General Variations

Plastomes of the three species of *Ludwigia* sp., *Lgh*, *Lpm* and *Lo,* are circular double-stranded DNA molecules (Figure 3) which are all (as shown in Table 1) approximately the same size: *Lo* is 159,396 bp long, making it the smallest, while *Lgh* is the largest with 159,584 bp, and *Lpm* is intermediate at 159,537 bp. The overall GC content is almost the same for the three species (37.4% for *Lo*, 37.3 % for *Lgh* and *Lpm*) and the GC contents of the IR regions are higher than those of the LSC and SSC regions (approximately 43.5 % compared to 35% and *ca.*32% respectively). Between the three species, the lengths of the total chloroplasts, LSC, SSC, and IR are broadly similar (approximately 90.2 kb for LSC, 19.8 kb for SSC and 24.8 kb for IB, see details Table 1) and the three plastomes are perfectly syntenic if we orient the SSC fragments the same way.

**Table 1.** The general characteristics of the 3 *Ludwigia* plastomes.

|  | *L.octovalvis* * | *L.grandiflora* | *L.peplodes* |
| --- | --- | --- | --- |
| **Size (bp)** |  |  |  |
|  | 159,396 | 159,584 | 159,537 |
| LSC | 90,183 | 90,272 | 90,156 |
| SSC | 19,703 | 19,788 | 19,799 |
| IR | 24,755 | 24,762 | 24,791 |
| **GC%** |  |  |  |
|  | 37.4 | 37.3 | 37.3 |
| LSC | 35.2 | 35.1 | 35.1 |
| SSC | 32 | 31.7 | 31.7 |
| IR | 43.5 | 43.5 | 43.4 |

\*KX827312 (reference)

All three *Ludwigia* sp. plastomes contain the same number of functional genes (134 in total) encoding 85 proteins (embracing 7 duplicated in the IR region: *ndh*B, *rpl2*, *rpl23*, *rps7*, *rps12*, *ycf2*, *ycf15*), 37 tRNAs (including *trn*K-UUU which contains *mat*K), and 8 rRNAs (16S, 23S, 5S, and 4.5S as duplicated sets in the IR). Among these genes, 18 contain introns, of which six are tRNAs (Table 2). Only the *rps12* gene is a trans-spliced gene. A total of 46 genes are involved in photosynthesis, and 71 genes related to transcription and translation, including a bacterial-like RNA polymerase and 70S ribosome, as well as a full set of transfer RNAs (tRNAs) and ribosomal RNAs (rRNAs). Six other protein-coding genes are involved in essential functions, such as *acc*D, which encodes the β-carboxyl transferase subunit of acetyl-CoA carboxylase, an important enzyme for fatty acid synthesis; *mat*K encodes for maturase K, which is involved in the splicing of group II introns; *cem*A, a protein located in the membrane envelope of the chloroplast is involved in the extrusion of protons and thereby indirectly allows the absorption of inorganic CO2 in the plastids; *clp*P1 which is involved in proteolysis, and; *ycf1*, *ycf2*, two ATPases members of the TIC translocon. Finally, a highly pseudogenized *ycf15* locus was annotated in the IR even though premature stop codons indicate loss of functionality.

**Table 2.** Genes present in the plastomes of *Ludwigia*.

| FUNCTION | NAME |
| --- | --- |
| **Photosynthesis** |  |
| Rubisco | *rbcL* |
| Photosystem I (PSI) | *psaA, psaB, psaC, psal, psaJ* |
| PSI assembly factors | *ycf3# (pafI), ycf4 (pafII)* |
| Photosystem II | *psbA, psbB, psbC, psbD, psbE, psbF, psbH, psbI, psbJ, psbK, psbL, psbM, pbf1 (psbN) psbT, psbZ* |
| ATP synthase | *atpA, atpB, atpE, atpF#, atpH, atpI* |
| Cytochrome *b6f* | *petA, petB#, petD#, petG, petL, petN* |
| Cytochrome biogenesis | *ccsA* |
| NADPH dehydrogenase | *ndhA#, ndhB\*\*#, ndhC, ndhD, ndhE, ndhF, ndhG, ndhH, ndhI, ndhJ* |
| **Transcription and translation** |  |
| Transcription | *rpoA, rpoB, rpoC1#, rpoC2* |
| Small ribosomal proteins | *rps2, rps3, rps4, rps7\*\*, rps8, rps11, rps12\*\*#, rps14, rps15, rps16#, rps18, rps19* |
| Large ribosomal proteins | *rpl2\*\*#, rpl14, rpl16#, rpl20, rpl22, rpl23\*\*, rpl32, rpl33, rpl36* |
| Translation initiation | *infA* |
| Ribosomal RNA | *rrn5\*\*, rrn4,5\*\*, rrn16\*\*, rrn23\*\** |
| Transfer RNA | *trnA-UGC\*\*#, trnC-GCA, trnD-GUC, trnE-UUC, trnF-GAA, trnFM-CAU, trnG-GCC, trnG-UCC#, trnH-GUG,, trnI-CAU\*\*, trnI-GAU\*\*#, trnK-UUU#, trnL-CAA\*\*, trnL-UAA#, trnL-UAG, trnM-CAU, trnN-GUU\*\*, trnP-UGG, trnQ-UUG, trnR-ACG\*\*, trnR-UCU, trnS-GCU, trnS-GGA, trnS-UGA, trnT-GGU, trnT-UGU, trnV-GAC\*\*, trnV-UAC#, trnW-CCA, trnY-GUA* |
| **Other functions** |  |
| Group II intron splicing | *matK* |
| Inorganic carbon uptake | *cemA* |
| Protease | *clpP1#* |
| Fatty acid synthesis/Heat tolerance | *accD* |
| TIC machinery (protein import) | *ycf1 (Tic214), ycf2\*\** |
| Unknown function pseudogene | *ycf15\*\** |

\*\* duplicated in IR region, # spliced genes

#### Segments Contractions/Expansion

The junctions between the different chloroplast segments were compared between three *Ludwigia* sp. (*Lpm*, *Lgh* and *Lo),* and we found that the overall resemblance of *Ludwigia* sp. plastomes was confirmed at all junctions (Figure 4A). In all three genomes, *rpl22, rps19, and rpl2* were located around the LSC/IRb border, and *rpl2*, *trn*H, and *psb*A were located at the IRa/LSC edge. The JSB (junction between IRb and SSC) is either located in the *ndh*F gene or the *ycf1* gene depending on the orientation of the SSC region (Figure 4B). The *ycf1* gene was initially annotated as a 1139 nt pseudogene that we biocurate as a larger gene (5302 nt) with a frameshift due to a base deletion, compared to *Lg* and *Lo* which both carry a complete *ycf1* gene.

**Figure 4:**
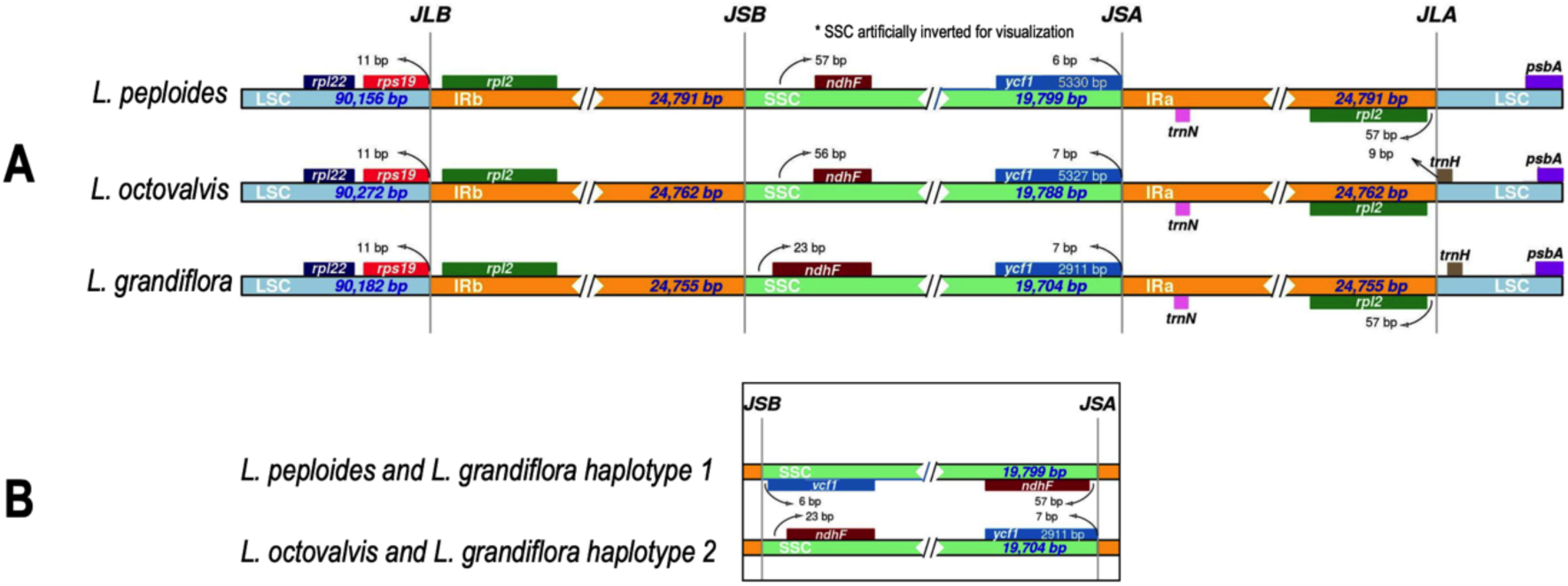
Comparison of the borders of LSC, SSC, and IR regions in Onograceae plastomes. **A:** Comparison of the junction between large single-copy (LSC, light blue), inverted repeat (IR, orange) and short single-copy (SSC, light green) regions among the chloroplast genomes of *L. octovalvis*, *L. peploides* and *L. grandiflora* (both haplotypes). Genes are denoted by colored boxes and the gaps between genes and boundaries are indicated by base lengths (bp). JLB: junction line between LSC and IRb; JSB: junction line between IRb and SSC; JSA: junction line between SSC and IRa; JLA: junction line between IRa and LSC. **B:** Comparison of SSC boundaries in haplotype 1 (*L. peploides* and *L. grandiflora* haplotype 1) and haplotype 2 (*L. octovalvis* and *L. grandiflora* haplotype 2) plastomes.

If we compare *Ludwigia* sp. chloroplastic LSC/SCC/IR junctions (via IRscope) with representative Onagraceae plastomes of *Chamaenerion sp. conspersum* (MZ353638) and *sp. angustifolium* (NC_052848), *Circaea sp. cordata* (NC_060876) and *sp. alpina* (NC_061010), *Epilobium amurense* (NC_061015) and *Oenothera villosa* subsp. *strigosa* (NC_061365) and *Oenothera lindheimeri* (MW538951) (Figure 5), we can observe that the gene positions at the JLB (junction of LSC/IRb) and JLA (junction of IRa/LSC) boundary regions are well-preserved throughout the entire family, whereas those at the JSB and JSA regions differ. Concerning JSB (junction of IRb/SSC), in the five Onagraceae genera studied, *ndh*F is duplicated, with the exception of *Circaea* sp. and *Ludwigia* sp. For *Oenothera villosa*, the first copy of *ndh*F, which is located in the IRb, overlaps the JSB border, whereas for *Oenothera lindheimeri*, *Epibolium amurense* and Chamaenerion sp., *ndh*F is only located in inverted repeats. Only *Circaea* sp. and *Ludwigia* sp. have a unique copy of this locus, and it is found in the SSC segment (Figure 5). At the JSA border (junction of SSC/Ira), in *Circaea* sp., the *ycf1* gene crosses the IRa/SSC boundary and extends into the IRa region.

**Figure 5:**
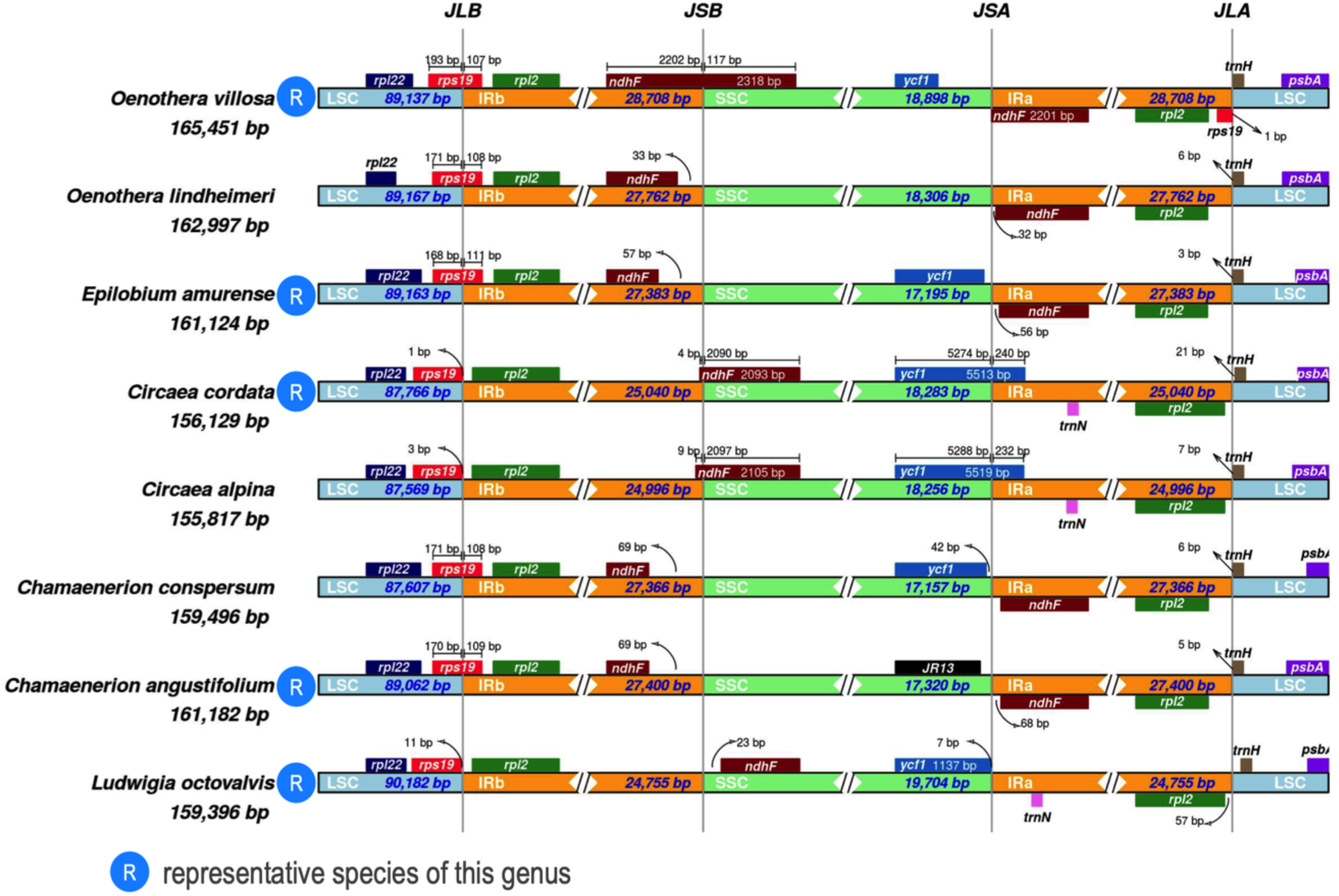
Comparison of LSC, SSC and IR regions boundaries in Onograceae chloroplast genomes. Representative sequences from each genus have been chosen (noted R on the diagram) except for *Oenothera lindheimeri* (only 89.35 % identity with others *Oenothera*), *Circaea alpina* (99.5 % identity but all others *Circaea* are 99.9% identical) and *Chamaenerion conspersum* (99% but all others *Chamaenerion* are ca. 99.7 identical). As shown in Figure 7, the 3 *Ludwigia* plastomas had the same structure*, L. octovalvis* was chosen as a representative of this genus. JLB: junction of LSC/IRb; JSB: junction of IRb/SSC; JSA: junction of SSC/IRa; JLA: junction of IRa/LSC. Accession numbers : *Chamaenerion sp. conspersum* (MZ353638), *Chamaenerion sp. angustifolium* (NC_052848), *Circaea sp. cordata* (NC_060876), *Circaea sp. alpina* (NC_061010), *Epilobium amurense* (NC_061015), *Oenothera villosa* subsp. *strigosa* (NC_061365) and *Oenothera lindheimeri* (MW538951).

When comparing the respective sizes of chloroplast fragments (IR/SSC/LSC) in Onagraceae, it can be observed that *Ludwigia* species exhibit expansions in the SSC and LSC regions which are not compensated by significant contractions in the IR regions. This is likely due to the relocation of the *ndh*F in the SSC region and *rps19* in the LSC region. Additionally, there may be significant size variations in the intergenic region between *trnI* and *ycf2*, as well as the intergenic segment containing the *ycf15* pseudogene (Add. Figure 4).

#### Repeats and SSRs analysis

In this study, we analyzed the nature and distribution of single sequence repeats (SSR), as their polymorphism is an interesting indicator in phylogenetic analyses. A total of 65 (*Lgh*), 48 (*Lpm*) and 45 (*Lo*) SSRs were detected, the majority being single nucleotide repeats (38–21), followed by tetranucleotides (12–10) and then di-, tri- and penta-nucleotides (Add. Figure 5A). Mononucleotide SSRs are exclusively composed of A and T, indicating a bias towards the use of the A/T bases, which is confirmed for all SSRs (Add. Figure 5B). In addition, the SSRs are mainly distributed in the LSC region for the three species, which is probably biased by the fact that LSC is the longest segment of the plastome (Add. Figure 5C). The analysis of SRR locations revealed that most were distributed in non-coding regions (intergenic regions and introns, Add. Figure 5D).

The chloroplast genomes of the three *Ludwigia* species were also screened for long repeat sequences. They were counted in a non-redundant way (if smaller repetitions were included in large repeats, only the large ones were considered). Four types of repeats (tandem, palindromic inverted and direct) were surveyed in the three *Ludwigia* sp. plastomes. No inverted repeats were detected with the criteria used.

For the three other types of repeats, here are their distributions:

*Tandem repeats* (Table 3A): Perfect tandem repeats (TRs) with more than 15 bp were examined. Twenty-two *loci* were identified in the three *Ludwigia* sp. plastomes (*Lgh*, *Lpm*, *Lo*), heterogeneously distributed as shown in Table 3A: 13 loci (plus one imperfect) in *Lo*, nine loci (plus one imperfect) in *Lgh* and seven loci (plus two imperfect) in *Lpm*. It can therefore be seen that the TR distributions (occurrence and location) are specific to each plastome, since only four pairs are common to the three species. Thus, nine TRs are unique to *Lo*, three to *Lpm* and three to *Lgh.* Two pairs are common to *Lgh* and *Lpm* and one is common to *Lo* and *Lgh.* TRs are mainly intergenic or intronic but are detected in two genes (*acc*D and *ycf1)*. These genes have accelerated substitution rates, although this does not generate a large difference in their lengths. This point will be developed later in this article.

**Table 3A:**
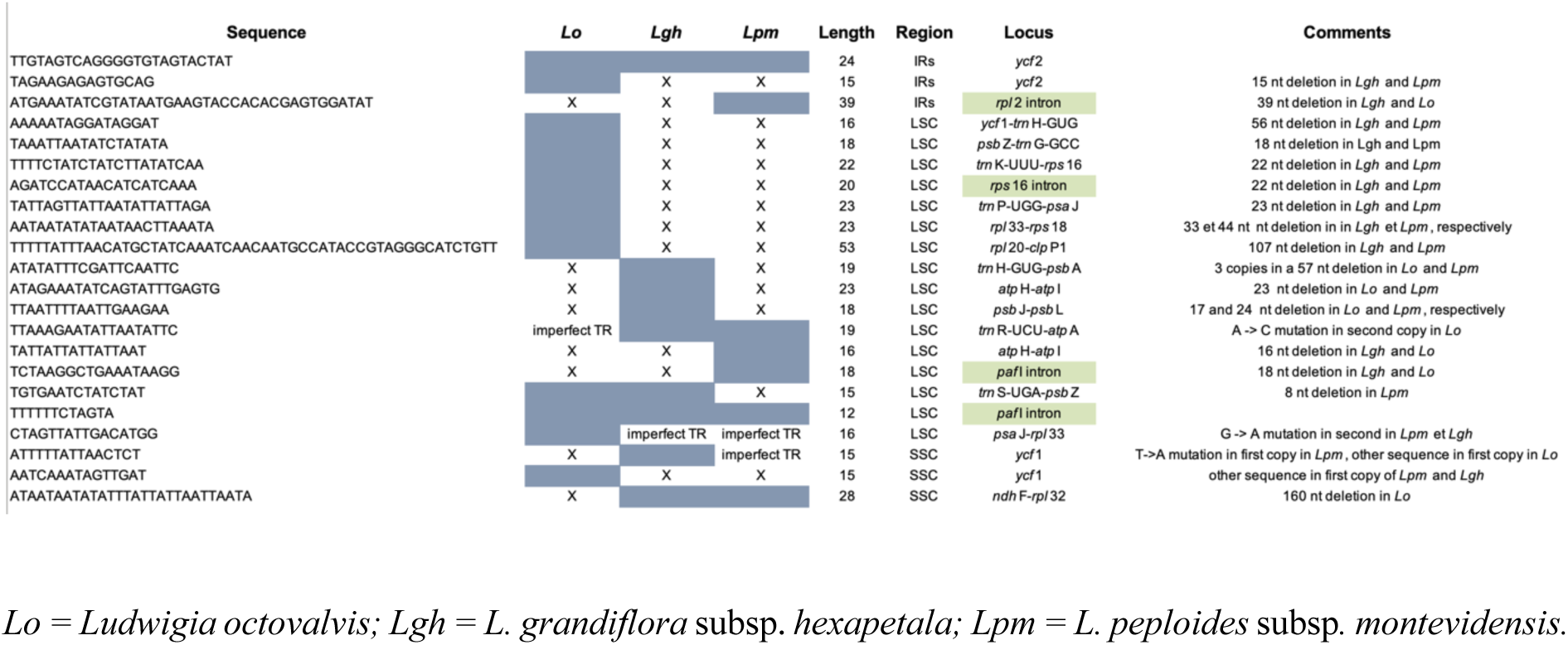
Tandem repeats.

*Direct repeats* (Table 3B): There are few direct (non-tandem) repeats (DRs) in the chloroplast genomes of *Ludwigia* sp. A single direct repeat of 41 nt is common to the three species, at 2 kb intervals, in *psaB* and *psaA* genes. This DR corresponds to an amino acid repeat [WLTDIAHHHLAIA] which corresponds to a region predicted as transmembrane. We then observe three direct repeats conserved in *Lpm* and *Lgh* in *ycf1*, *acc*D and *clp*P1 respectively, two unique DRs in *Lo* (in the *acc*D gene and *rps12-clp*P1 intergene) and one in *Lgh* (in the *clp*P1 intron 1 and *clp*P1 intron 2).

**Table 3B:**
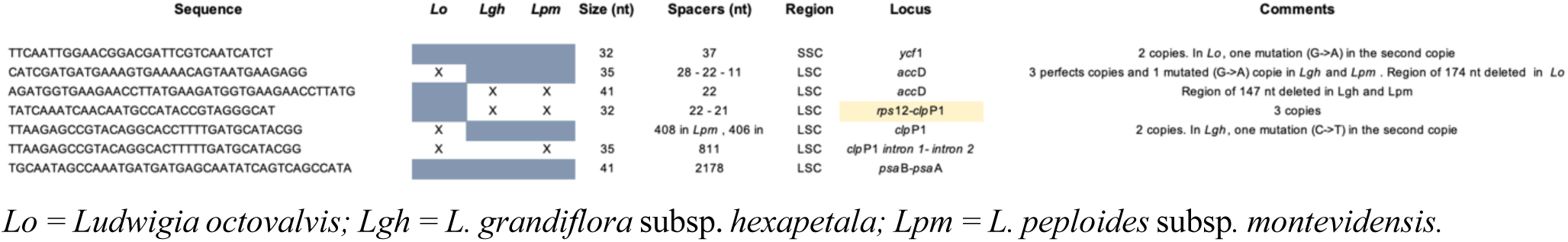
Direct repeats.

*Palindromes* (Table 3C): Palindromic repeats make up the majority of long repetitions, with the numbers of perfect repeats varying from 19, 24 and 26 in *Lo*, *Lgh* and *Lpm*, respectively, and the number of quasi-palindromes (1 mutation) varying between 8, 3 and 6. They are mainly found in the intronic and intergenic regions, with the exception of six genic locations in *psb*D, *ndh*K, *ccs*A and *rpl22,* and two palindromic sequences in *ycf2.* These gene palindromic repeats do not seem to cause genetic polymorphism in *Ludwigia* and can be considered as silent.

**Table 3C:**
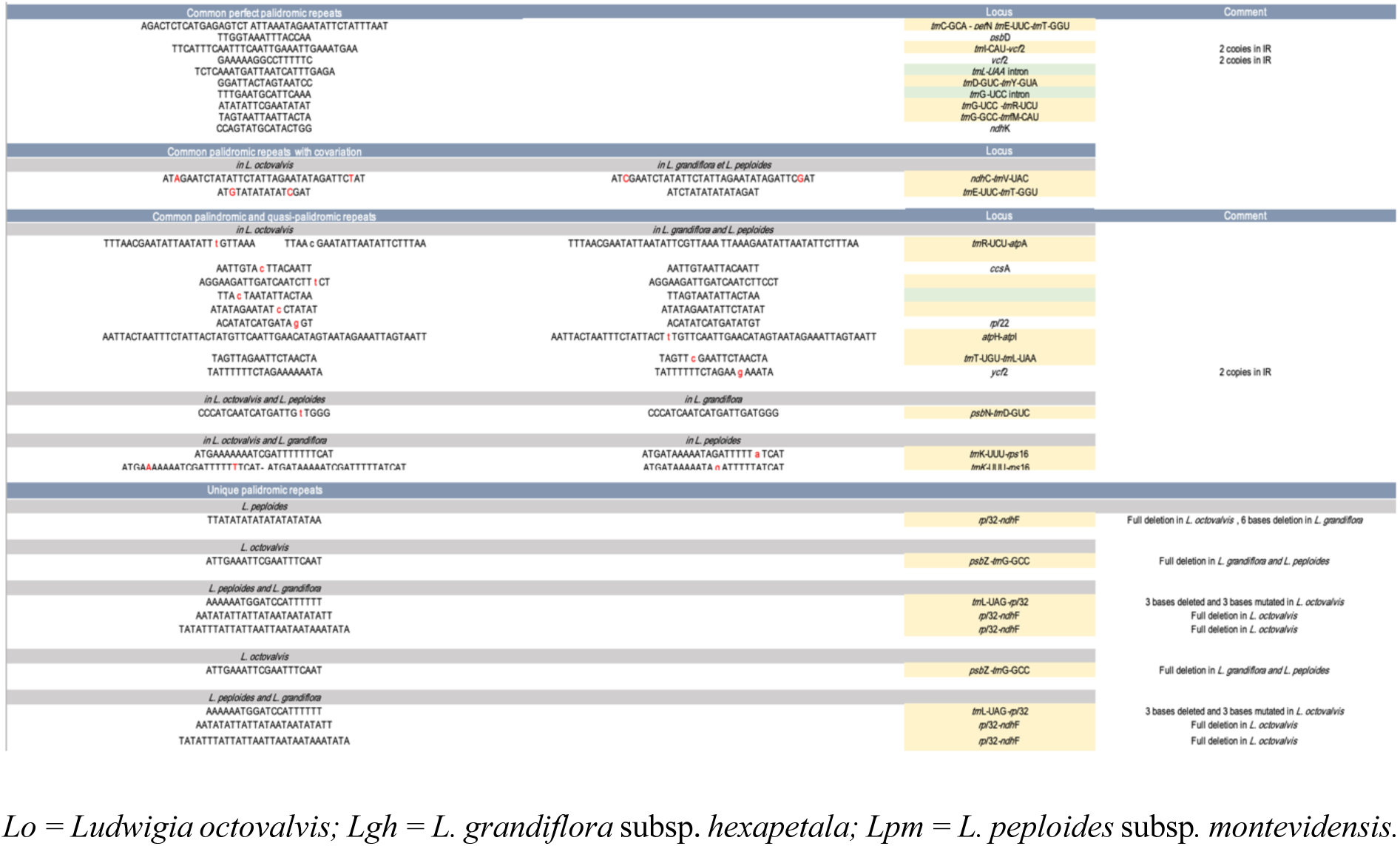
Palindromic repeats.

Thirteen palindromes are common to the three species (including 2 with co-variations in *Lo*). 13 others present in *Lpm* and *Lgh* correspond to quasi-palindromes (QPs) in *Lo* due to mutated bases, and conversely, three *Lo* perfect palidromes are mutated in *Lpm* and *Lgh*. Finally, only five palindromes are species specific. Two in particular are located in the hypervariable intergenic spacer *ndh*F*-rpl32*, and are absent in *Lo* due to a large deletion of 160 nt.

#### Repeat distribution in LSC, SSC and IR segments

In the IRa/IRb regions, repeats are only identified in the first 9 kb region between *rpl2* and *ycf2*: a tandem repeat in the *Lpm rpl2* intron, and a tetranucleotide repeat, [TATC]*3, located in the *ycf2* gene in the 3 species. In *ycf2* we also found 1 common palindrome (16 nt), a single palindrome in *Lo* (20 nt, absent following an A:G mutation in the 2 other species), as well as a shared tandem repeat (24 nt), and an additional 15 nt tandem repeat in *Lo* which adds 4 amino acids to protein sequence.

In the SSC region, the repeats are almost all located in the intergenic and/or intronic regions, with a hotspot between *ndh*F and *ccs*A. There is also a shared microsatellite in *ndh*F, and a palidrome (16 nt) in *ccs*A which is absent in *Lo* (due to an A:C mutation), resulting in a synonymous mutation (from isoleucine to leucine). We also observed multiple and various repeats in the *ycf1* gene: 3 common poly-A repeats (from 10 to 13 nt), 3 species-specific microsatellites (ATAG)*3 and (ACCA)*4 in *Lgh* and (CAAC)*3 in *Lo,* as well as two direct repeats of 32 nt (37 nt spacing), which were absent from *Lo* due to a G:T SNP. Two tandem repeats were also observed in *Lo* and *Lgh*. Neither of these repeats are at the origin of the frameshift causing the pseudogenization of *ycf1* in *Lo*, this latter being due to a single deletion of an A at position 3444 of the gene.

Finally, in the LSC region, the longest segment, which consequently contains the maximum number of repeats, we still observed a preferential localization in the intergenic and intronic regions since only genes *atp*A, *rpo*C2, *rpo*B, *psb*D, *psb*A, *psb*B, *ndh*K and *clp*P1 contain either mononucleotic repeats (poly A and T), palindromes, or microsatellites (most often common to the three species and without affecting the sequences of the proteins produced). As mentioned earlier, the only exception is the *acc*D gene, which contains several direct and tandem repeats in *Lgh* and *Lpm,* corresponding to a region of 174 nt (58 amino acids) missing in *Lo* and, conversely, a direct repeat of 40 nucleotides, in a region of 147 nt (49 aa), which is present in *Lo* and missing in the other two species. These tandem repeats lead to the presence of four copies of 9 amino acids [DESENSNEE] in *Lgh* and *Lpm*, two of which form a larger duplication of 17 aa [FLSDSDIDDESENSNEE]. Similarly, the TRs present only in *Lo* generate two perfect 9 amino acid repeats [EELSEDGEE], included in two longer degenerate repeats of 27 nt (Add. Figure 6). It should be noted that though these TRs do not disturb the open reading phases, it is still possible for them to form an intron which is not translated. Different functional studies will be necessary to clarify this point. The presence of polymorphisms of the *acc*D gene between *Lo* and the two species (*Lpm*, *Lgh*) is interesting because *acc*D, that encodes a subunit of acetyl-CoA carboxylase (EC 6.4.1.2). This enzyme is essential in fatty acid synthesis and also catalyzes the synthesis of malonyl-CoA, which is necessary for the growth of dicots, plant fitness and leaf longevity, and is involved in the adaptation to specific ecological niches (Konishi and Sasaki 1994). Large *acc*D expansions due to TRs have also been described in other plants such as *Medicago* (S. Wu et al. 2021) and *Cupressophytes* (J. Li, Su, and Wang 2018). Some authors have suggested that these inserted repeats are not important for acetyl-CoA carboxylase activity as the reading frame is always preserved, and they assume that these repeats must have a regulatory role (Gurdon and Maliga 2014).

#### Sequence Divergence Analysis and Polymorphic Loci Identification

Determination of divergent regions by MVista, using *Lo* as a reference, confirmed that the three *Ludwigia* sp. plastomes are well preserved if the SSC segment is oriented in the same way (Add. Figure 7). Sliding window analysis (Figure 6) indicated variations in definite coding regions, notably *clp*P, *acc*D, *ndh5*, *ycf1* with high Pi values, and to a lesser extent, *rps16*, *mat*K, *ndh*K, *pet*A, *ccs*A and four tRNAs (*trn*H, *trn*D, *trn*T and *trn*N). These polymorphic *loci* could be suitable for inferring genetic diversities in *Ludwigia* sp.

**Figure 6:**
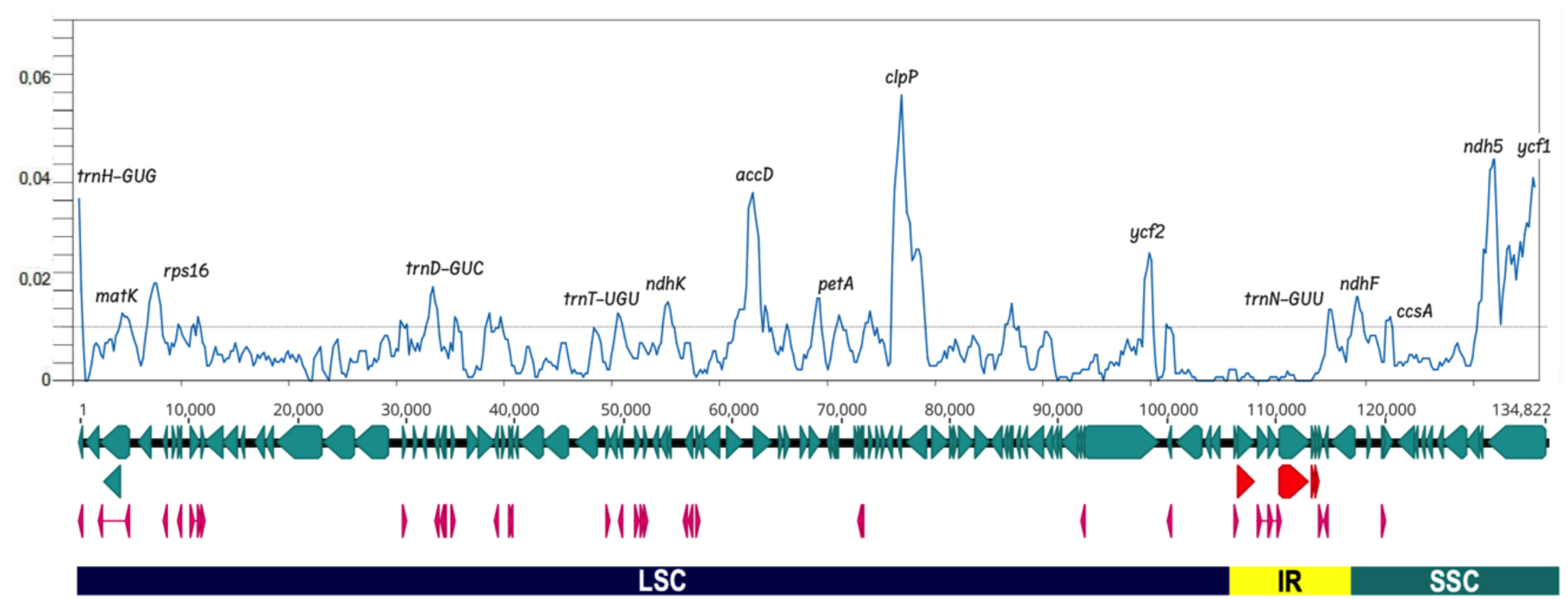
Illustration of nucleotide diversity of the three *Ludwigia* chloroplast genome sequences. The graph was generated using DnaSP software version 6.0 (windows length: 800 bp, step size: 200 bp) (Julio Rozas et al. 2017)(J. Rozas and Rozas 1999). The x-axis corresponds to the base sequence of the alignment, and the y-axis represents the nucleotide diversity (π value). LSC, SSC and IR segments were indicated under the line representing the genes coding the proteins (in light blue) the tRNAs (in pink) and the rRNAs (in red). The genes marking diversity hotspots are noted at the top of the peaks.

A comparative analysis of the sizes of protein coding genes sizes also shows that the *rps11* gene initially annotated in *Lo* is shorter than those which have been newly annotated in *Lgh* and *Lpm* (345 bp instead of 417 bp). Comparative analysis by BLAST shows that it is the long form which is annotated in other Myrtales, and the observation of the locus in *Lo* shows a frameshift mutation (deletion of a nucleotide in position 311). Functional analysis would be necessary to check whether the *rps11* frameshift mutation produces shorter proteins that have lost their function. And only obtaining the complete genome will verify whether copies of some of these genes have been transferred to mitochondrial or nuclear genomes. Such *rps11* horizontal transfers have been reported for this gene in the mitochondrial genomes of various plant families(Richardson and Palmer 2007). This also applies to *ycf1*, found as a pseudogene in *Lo* (as specified previously), although it is not known if this reflects a gene transfer or a complete loss of function (de Vries et al. 2015)(Filip and Skuza 2021). Moreover, there is a deletion of nine nucleotides in the 3’ region of the *rpl32* gene in *Lgh* and *Lpm*, leading to a premature end of the translation and the deletion of the last 4 amino acids [QRLD], which are replaced by a K. However, if we look carefully at the preserved region as defined by the RPL32 domain (CHL00152, member of the superfamily CL09115), we see that the later amino acids are not important for *rpl32* function since they are not found in the orthologs.

Our results show that the Ka/Ks ratio is less than 1 for most genes (Figure 7). This indicates adaptive pressures to maintain the protein sequence except for *mat*K (1.17 between *Lgh* and *Lpm*), *acc*D (2.48 between *Lgh* and *Lo* and 2.16 between *Lpm* and *Lo*), *ycf2* (4.3 between both *Lgh-Lp* and *Lo*) and *ccs*A (1.4 between both *Lgh-Lpm* and *Lo*), showing a positive selection for these genes, and a possible key role in the processes of the species’ ecological adaptations. As we have already described the variability in the *acc*D sequence, we will focus on *ycf2*, *mat*K, and *ccs*A variations.

**Figure 7:**
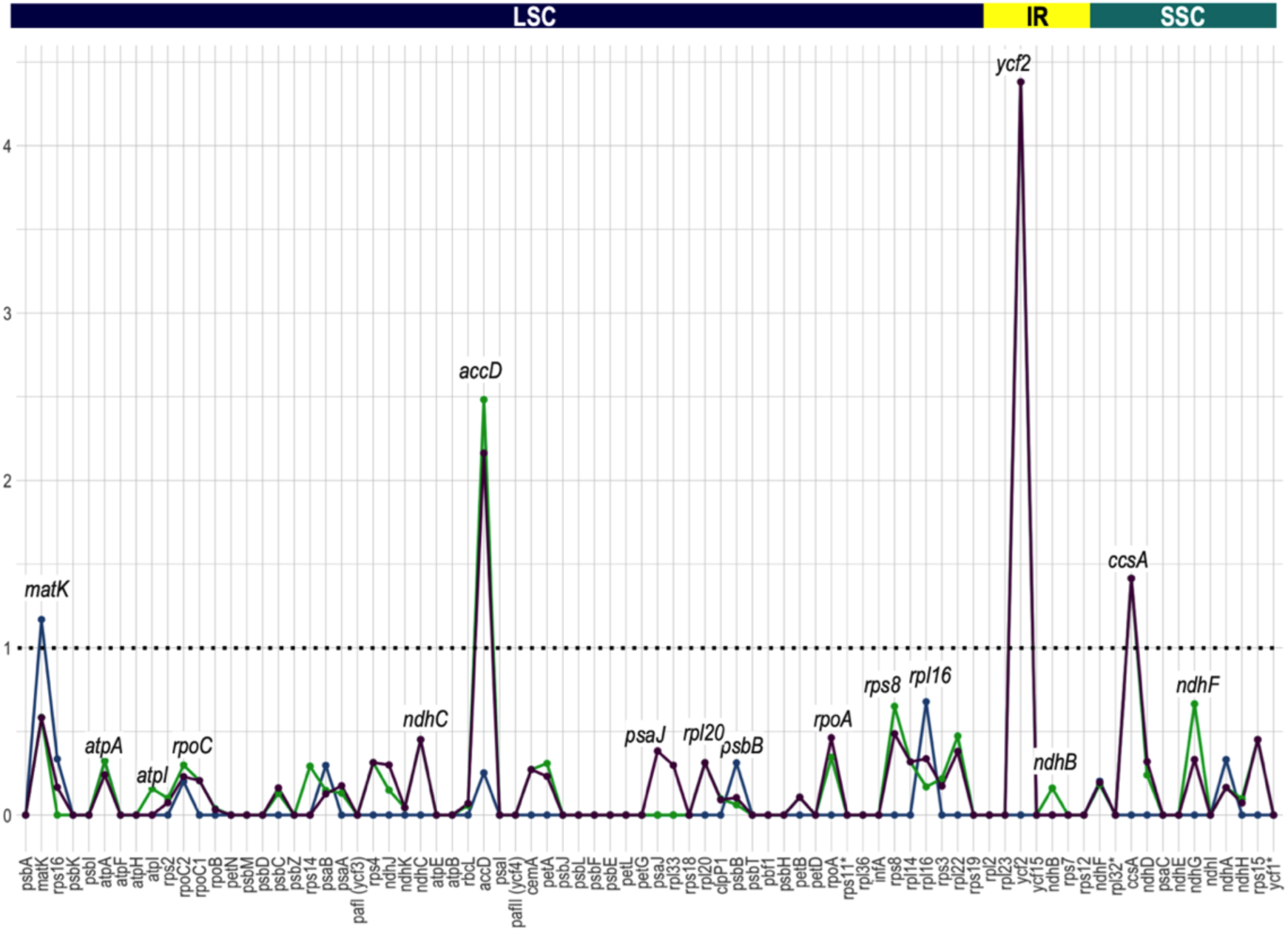
The Ka/Ks ratios of the 80 protein-coding genes of *Ludwigia* plastomes. The blue curve represents *L. grandiflora* versus *L. peploides*, purple curve denotes *L. grandiflora* versus *L. octovalvis* and green curve *L. peploides* versus *L. octovalvis.* Four genes (*matK*, *accD*, *ycf2* and *ccsA*) have Ka/Ks ratios greater than 1.0, whereas the Ka/Ks ratios of the other genes were less than 1.0.

Concerning *ccs*A, the variations observed, although significant, concern only five amino acids, and modifications do not seem to affect the C-type cytochrome synthase gene function.

Concerning *ycf2*, our analysis shows that this gene is highly polymorphic with 256 SNPs that provoke 10 deletions, 7 insertions, 21 conservative and 49 non-conservative substitutions in *Lo* (Add. Figure 8), compared to *Lgh* and *Lpm* (100 % identical). This gene has been shown as “variant” in other plant species such as *Helianthus tuberosus* (Q. Zhong et al. 2019).

The *mat*K gene has been used as a universal barcoding locus to enable species discrimination of terrestrial plants (Antil et al. 2023), and is often, together with the *rbc*L gene, the only known genetic resource for many plants. Thus, we propose a phylogenetic tree from *Ludwigia mat*K sequences (Figure 8). It should however be noted that this tree contains only 149 amino acids common to all the sequences (out of the 499 in the complete protein). As only three complete *Ludwigia* plastomes are available at the time of our study, we cannot specify whether these barcodes are faithful to the phylogenomic history of *Ludwigia* in the same way as the complete plastome. In any case, for this tree, we can see that *Lo* stands apart from the other *Ludwigia* sp., *Lpm* and *Lgh*, and that the *L. grandiflora* subsp. *hexapetala* belongs to the same branch as the species *L. ovalis* (aquatic taxon used in aquariums (J. Li et al. 2022)), *L. stolonifera* (native to the Nile, found in a variety of habitats, from freshwater wetlands to brackish and marine waters) (Soliman, Hamdy, and Hamed 2018) and *L. adscendens* (common weed of rice fields in Asia) (Kamoshita et al. 2016). *Lpm* is in a sister branch, close to the *L. grandiflora* subsp. *hexapetala*, forming a phylogenetic group corresponding to subsect Jussiaea (in green, Figure 8).

**Figure 8:**
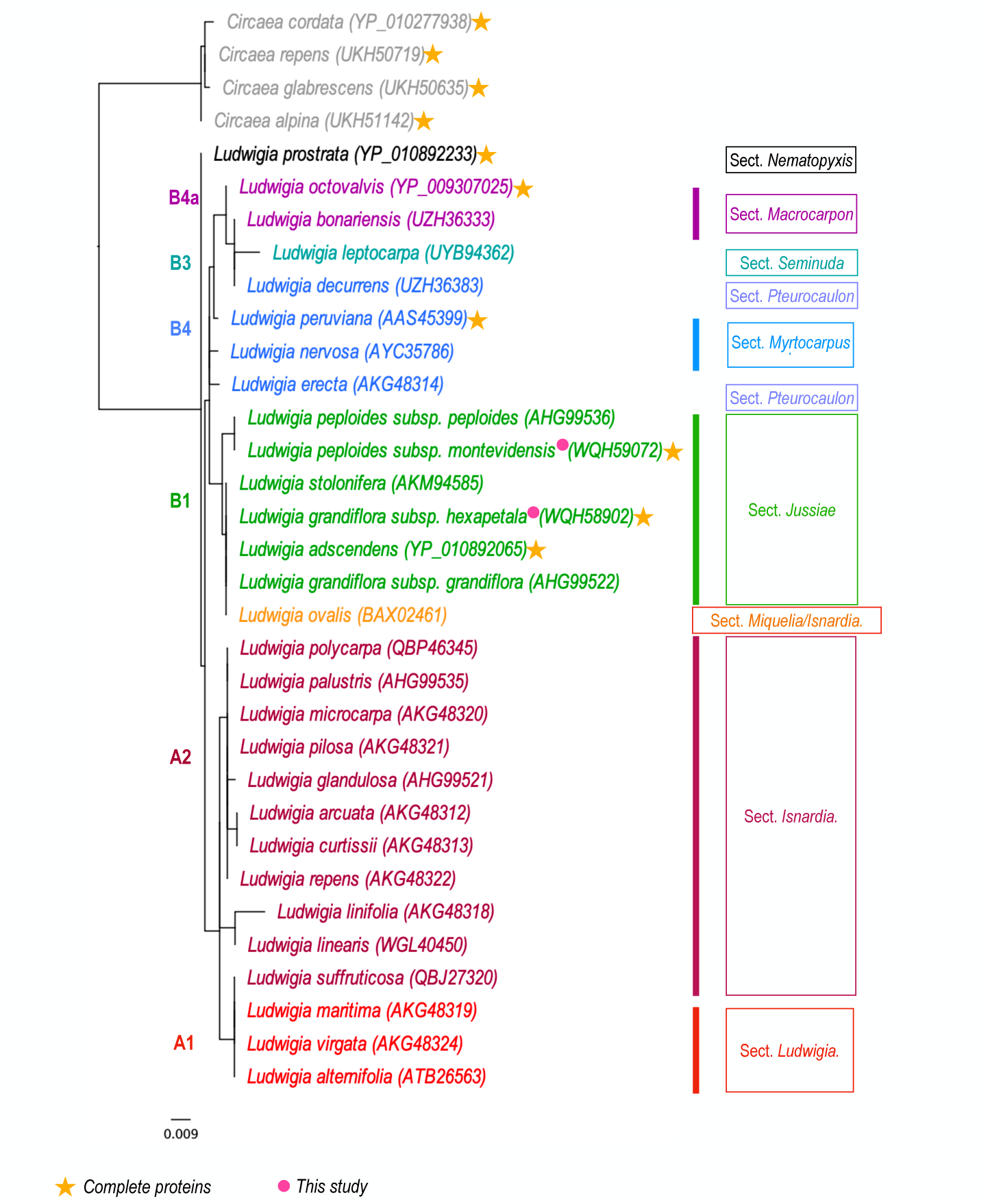
Phylogenetic tree based on *Ludwigia* MatK protein sequences. Only six *Ludwigia* sequences are complete (yellow star), the others correspond to amino acids ranging from 128 to 289 aa, with an average of 244 aa. Clades are named and colored regarding the *Ludwigia* phylogeny proposed by Liu et al. (2017) (S. H. Liu et al. 2017). The sections are based on the works of Raven (1963) (Raven P.H. 1963), Wagner et al (2017) (Wagner, Hoch, and Raven 2007b)and Liu et al. (2023) (S. H. Liu et al. 2023). The scale bar indicates the branch length.

## Discussion

In the present study, we first sequenced and de novo assembled the chloroplast (cp) genomes of Ludwigia peploides (Lpm) and Ludwigia grandiflora (Lgh), two species belonging to the Onagraceae family. We employed a hybrid strategy and demonstrated the presence of two cp haplotypes in Lgh and one haplotype in Lpm, although the presence of both haplotypes in Lpm is likely. Furthermore, we compared these genomes with those of other species in the Onagraceae family to expand our knowledge of genome organization and molecular evolution in these species.

Our findings demonstrate that the utilization of solely short reads has failed to produce complete Ludwigia plastomes, likely due to challenges posed by long repeats and rearrangements. On the other hand, relying solely on long reads resulted in a lower quality sequence due to insufficient coverage and sequencing errors. After conducting our research, we discovered that, for Lgh plastomes, hybrid assembly, which incorporates both long and short read sequences, resulted in the most superior complete assemblies. This innovative approach capitalizes on the advantages of both sequencing technologies, harnessing the accuracy of short read sequences and the length of long read sequences. In the case of our study on Lgh plastome reconstruction, hybrid assembly was the most complete and effective, similarly to studies on other chloroplasts, such as those in Eucalyptus (Wang et al. 2018b), Falcataria (Anita, Matra, and Siregar 2023), Carex (S. Xu et al. 2023) or Cypripedium (Guo et al. 2021).

In our study, we were able to identify the presence of two haplotypes in Lgh, which is a first for Ludwigia (and more broadly within Onagraceae), as the plastome of L. octovalvis was only delivered in one haplotype (S.-H. Liu et al. 2016).

Due to the unavailability of sequence data for Ludwigia octovalvis and the fact that we only have long reads for Ludwigia peploides, none of which large enough to cover the SSC/IR junctions, we are unable to conclusively identify the presence of these two forms in the Ludwigia genus. However, we believe that they are likely to be present. Unfortunately, the current representation of plastomes in GenBank primarily consists of short-read data, which may result in an underrepresentation of this polymorphism. It is unfortunate that structural heteroplasmy, which is expected to be widespread in angiosperms, has been overlooked. Existence of two plastome haplotypes has been identified in the related order of Myrtales (Eucalyptus sp.), in 58 species of Angiosperms, (Wang and Lanfear 2019), Asparagales (Ophrys apifera orchid (Bateman et al. 2021)), Brassicales (Carica papaya, Vasconcellea pubescens (Lin et al. 2020)), Solanales (Solanum tuberosum (Lihodeevskiy and Shanina 2022)), Laurales (Avocado Persea americana (Nath et al. 2022)) and Rhamnaceae (Rhamnus crenata (Wanichthanarak et al. 2023)). However, the majority of reference plastomes in the current GenBank database (Release 260: April 15, 2024) are described as a single haplotype, indicating an underrepresentation of structural heteroplasmy in angiosperm chloroplasts. This underscores the importance of sequencing techniques, as the database is predominantly composed of short-read data (98%), which are less effective than long reads or hybrid assemblies at detecting flip-flop phenomena in the LSC region.

The chloroplast genome sizes for the three genera of Onagraceae subfam. Onagroideae varied as follows: Circaea sp. ranged from 155,817 bp to 156,024 bp, Chamaenerion sp. ranged from 159,496 bp to 160,416 bp, and Epilobium sp. ranged from 160,748 bp to 161,144 bp (Luo et al. 2021). Our study revealed that the size of the complete chloroplast of Ludwigia (Onagraceae subfamily Ludwigioideae) ranged from 159,369 bp to 159,584 bp, which is remarkably similar to other Onagraceae plants (average length of 162,030 bp). Furthermore, Ludwigia plastome sizes are consistent with the range observed in Myrtales (between 152,214 to 171,315 bp (X. F. Zhang et al. 2021)). In the same way, similar overall GC content was found in Ludwigia sp. (from 37.3 to 37.4%), Circaea sp. (37.7 to 37.8%), Chamaenerion sp. and Epilobium sp. (38.1 to 38.2%,(Luo et al. 2021)) and more generally for the order Myrtales (36.9–38.9%, with the average GC content being 37%,(X. F. Zhang et al. 2021)). Higher GC content of the IR regions (43.5%) found in Ludwigia sp. has already been shown in the Myrtales order (39.7–43.5%) and in other families/orders such as Amaranthaceae (order Caryophyllales (J. Xu et al. 2020)) or Lamiaceae (order Lamiales (Lian et al. 2022)), and is mainly due to the presence of the four GC rich rRNA genes.

The complete chloroplast genomes of the three Ludwigia species encoded an identical set of 134 genes including 85 protein-coding genes, 37 tRNA genes and eight ribosomal RNAs, consistent with gene content found in the Myrtales order, with a gene number varying from 123 to 133 genes with 77–81 protein-coding genes, 29–31 tRNA gene and four rRNA genes (X. F. Zhang et al. 2021). Chloroplast genes have been selected during evolution due to their functional importance(Mohanta et al. 2020). In our current study, we made the noteworthy discovery that matK, accD, ycf2, and ccsA genes were subjected to positive selection pressure. These genes have frequently been reported in literature as being associated with positive selection, and are known to play crucial roles in plant development conditions. Lgh and Lpm are known to thrive in aquatic environments, where they grow alongside rooted emergent aquatic plants, with their leaves and stems partially submerged during growth, as reported by Wagner et al. in 2007 (Wagner, Hoch, and Raven 2007a). Both species possess the unique ability of vegetative reproduction, enabling them to establish themselves rapidly in diverse habitats, including terrestrial habitats, as noted by Haury et al (Haury et al. 2014b). Additionally, Lo is a wetland plant that typically grows in gullies and at the edges of ponds, as documented by Wagner et al. in 2007 (Wagner, Hoch, and Raven 2007a). Given their ability to adapt to different habitats, these species may have evolved specialized mechanisms to cope with various abiotic stresses, such as reduced carbon and oxygen availability or limited access to light in submerged or emergent conditions. Concerning matK, Barthet et al (Barthet and Hilu 2007) demonstrated the relationship between light and developmental stages, and MatK maturase activity, suggesting important functions in plant physiology. This gene has recently been largely reported to be under positive selection in an aquatic plant (Anubias sp.,(L. Li et al. 2022)), and more generally in terrestrial plants (Pinus sp (Zeb et al. 2022)or Chrysosplenium sp. (Z. Wu et al. 2020)). The accD gene has been described as an essential gene required for leaf development (Kode et al. 2005) and longevity in tobacco (Nicotiana tabacum)(Madoka et al. 2002). Under drought stress, plant resistance can be increased by inhibiting accD (Gu et al. 2020), and conversely, enhanced in response to flooding stress by upregulating accD accumulation (Bharadwaj et al. 2023). Hence, we can hypothesize that the positive selection observed on the accD gene can be explained by the submerged and emerged constraints undergone by Ludwigia species. The ycf2 gene seems to be subject to adaptive evolution in Ludwigia species. Its function, although still vague, would be to contribute to a protein complex generating ATP for the TIC machinery (proteins importing into the chloroplasts (Kikuchi et al. 2018)(Schreier et al. 2018)), as well as plant cell survival (Drescher et al. 2000)(Xing et al. 2022). The ccsA gene positive selection is found in some aquatic plants such as Anubia sp.(L. Li et al. 2022), marine flowering plants as Zostera species (J. Chen et al. 2023), and some species of Lythraceae (Gu et al. 2020). The ccsA gene is required for cytochrome c biogenesis (Xie and Merchant 1996) and this hemoprotein plays a key role in aerobic and anaerobic respiration, as well as photosynthesis (Kranz et al. 1998). Furthermore, we showed that Lgh colonization is supported by metabolic adjustments mobilizing glycolysis and fermentation pathways in terrestrial habitats, and the aminoacyl-tRNA biosynthesis pathway, which are key components of protein synthesis in aquatic habitats (K Billet et al. 2018). It can be assumed that the ability of Ludwigia to invade aquatic and wet environments, where the amount of oxygen and light can be variable, leads to a high selective pressure on genes involved in respiration and photosynthesis.

Molecular markers are often used to establish population genetic relationships through phylogenetic studies. Five chloroplasts (rps16, rpl16, trnL-trnF, trnL-CD, trnG) and two nuclear markers (ITS, waxy) were used in previous phylogeny studies of Ludwigia sp.(S.-H. Liu et al. 2017). However, no SSR markers had previously been made available for the Ludwigia genus, or more broadly, the Onagraceae. In this study, we identified 45 to 65 SSR markers depending on the Ludwigia species. Most of them were AT mononucleotides, as already recorded for other angiosperms (Maheswari, Kunhikannan, and Yasodha 2021)(Y. Zhang et al. 2016). In addition, we identified various genes with highly mutated regions that can also be used as SNP markers. Chloroplast SSRs (cpSSRs) represent potentially useful markers showing high levels of intraspecific variability due to the non-recombinant and uniparental inheritance of the plastomes (Huang et al. 2018)(Leontaritou et al. 2021). Chloroplast SSR characteristics for Ludwigia sp. (location, type of SSR) were similar to those described in most plants. While the usual molecular markers used for phylogenetic analysis are nuclear DNA markers, cpSSRs have also been used to explore cytoplasmic diversity in many studies (Snoussi et al. 2022)(Song et al. 2014)(Wheeler et al. 2014). To conclude, the 13 highly variable loci and cpSSRs identified in this study are potential markers for population genetics or phylogenetic studies of Ludwigia species, and more generally, Onagraceae.

Concerning the MatK-based phylogenetic tree, its topology is generally congruent with the first molecular classification of Liu et al. (S.-H. Liu et al. 2017)as all Ludwigia from sect Jussiaea (clade B1) and sect. Ludwigia (clade A1) and sect. Isnardia (clade A2) branched together. In this MatK-based tree, Ludwigia prostrata, a species absent from previously published phylogenetic studies, positions itself alone at the root of the Ludwigia tree. This species, sole member of section Nematopyxis, is related as having no close relatives (Raven and Tai 1979), finding supported by our work. We also observed that Ludwigia ovalis branches within sect. Jussiaea, as its 258 amino acids partial MatK sequence (ca. half of the complete sequence) is identical to the MatK proteins of L. grandiflora, L. stolonifera and L. adscendens. Its phylogenetic placement remains unresolved: classified alone by Raven (1963) (Raven P.H. 1963) and Wagner (2017) (Wagner, Hoch, and Raven 2007b) in sect. Miquelia, later positioned by Liu et al. (2017)(S. H. Liu et al. 2017) within the Isnardia-Microcarpium section (using nuclear DNA) or as sister to it (using plastid DNA). For this reason, conducting a whole plastome analysis would be valuable to provide insights into L. ovalis phylogenetic positioning. Another species positioned on the margins of sect. Isnardia (clade A2) is Ludwigia suffruticosa (previously classified in sect. Microcarpium), which branches within sect. Ludwigia (clade A1). This positioning raises questions about the current grouping of sections Isnardia, Michelia, and Microcarpium into a single section Isnardia as proposed by Liu et al. (2023) (S. H. Liu et al. 2023)and highlights that plastid protein coding markers can provide differing phylogenetic insights. Finally, the last species positioned differently of this clade (clade B4) is Ludwigia decurrens (sect. Pterocaulon) which clusters with L. leptocarpa (clade B3) and L. bonariensis (clade B4a). However, it is important to note that in their study, Liu et al. (2017) indicate that clade B4 is moderately supported and that the two members of sect. Pterocaulon, L. decurrens and L. nervosa, diverge in all trees (S. H. Liu et al. 2017). In summary, acquiring complete plastomes for Ludwigia sp. could significantly enhance our understanding of the phylogeny of this complex genus. Furthermore, comparing nuclear and plastid phylogenies would help determine if they reflect the same evolutionary history and whether plastid phylogeny alone can accurately reconstruct the phylogeny of Ludwigia genus.

## Conclusion

In this study, we conducted the first-time sequencing and assembly of the complete plastomes of *Lpm* and *Lgh*, which are the only available genomic resources for functional analysis in both species. We were able to identify the existence of two haplotypes in *Lgh*, but further investigations will be necessary to confirm their presence in *Lo* and *Lpm*, and more broadly, within the *Ludwigia* genus. Comparison of all 10 Onagraceae plastomes revealed a high degree of conservation in genome size, gene number, structure, and IR boundaries. However, to further elucidate the phylogenetic analysis and evolution in *Ludwigia* and Onagraceae, additional chloroplast genomes will be necessary, as highlighted in recent studies of Iris and Aristidoideae species (Feng et al. 2022).

## Declarations

### Availability of data and materials

The datasets generated and/or analysed during the current study were available in GenBank (for *Lgh* haplotype 1, (LGH1) OR166254 and *Lgh* haplotype 2, (LGH2) OR166255; for *Lpm* haplotype, (LPM) OR166256). Chloroplastic short and long reads are available at EBI-ENA database (https://www.ebi.ac.uk/ena/browser/home) under these accession numbers for LGH plastomes (Long reads : Experiment : ERX13439011 ; Run : ERR14035997 and short reads : Experiment : ERX13439002 ; Run : ERR14035988) and for LPM plastomes (Long reads : Experiment : ERX13439014 ; Run : ERR14036000).

### Conflict of interest disclosure

The authors declare that they comply with the PCI rule of having no financial conflicts of interest in relation to the content of the article

### Funding

The post-doctoral research grant of Anne-Laure Le Gac was supported by the Conseil regional Bretagne (SAD18001).

## Acknowledgements

We are grateful to Luis Portillo-Lemus for developing the high molecular weight genomic DNA extraction protocol. All sequencing experiments were performed at the PGTB (doi:10.15454/1.5572396583599417E12).

## Additional information

**Add. Figure 1:**
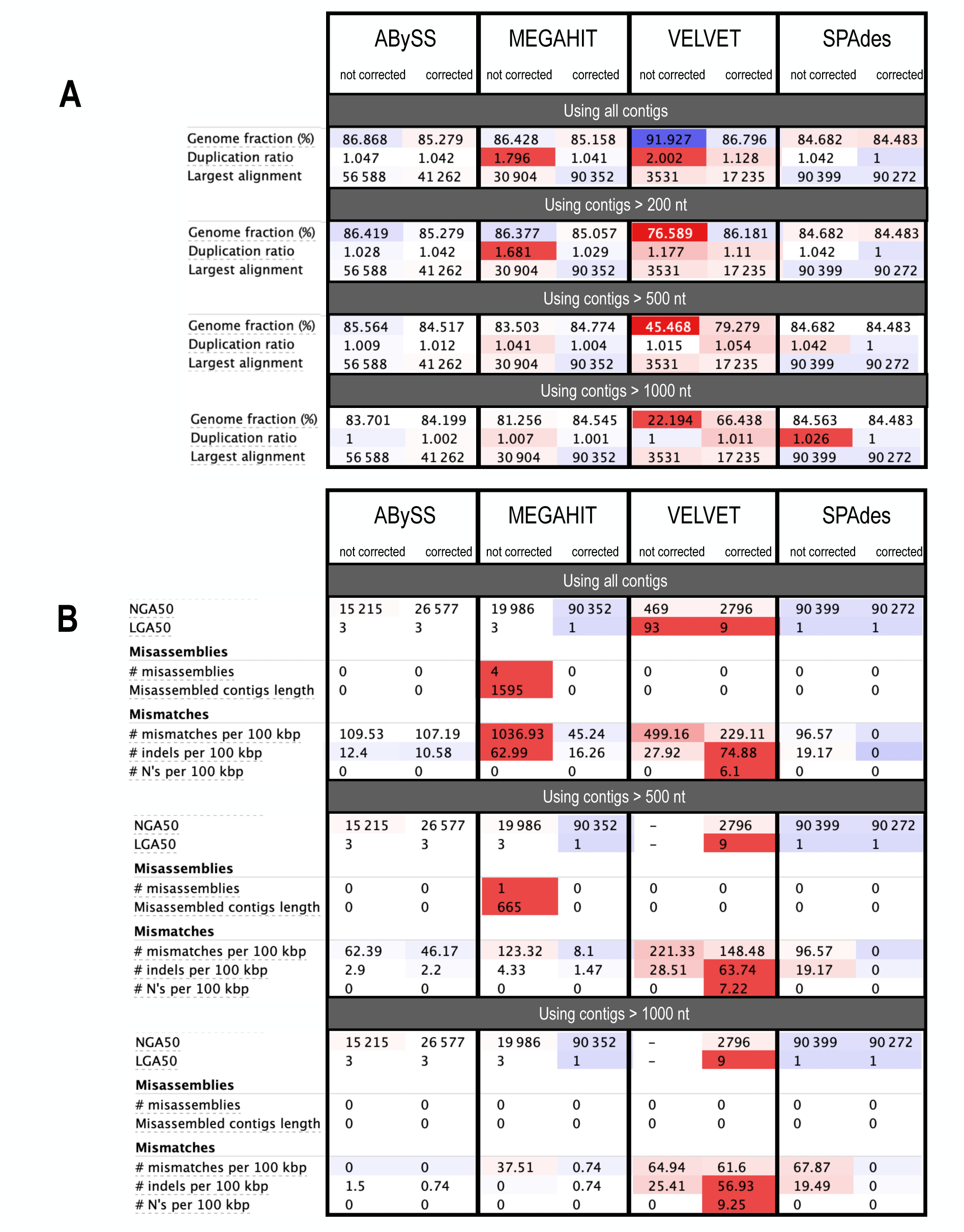
QUAST evaluation of performance of the four assembly tools (using corrected or uncorrected SRs). A: Comparison of plastome fraction, duplication rate and size of the largest alignment obtained. B: Comparison of classic metrics (NGA50 and LGA50), number of errors (misassemblies and mismatches) produced.

**Add. Figure 2:**
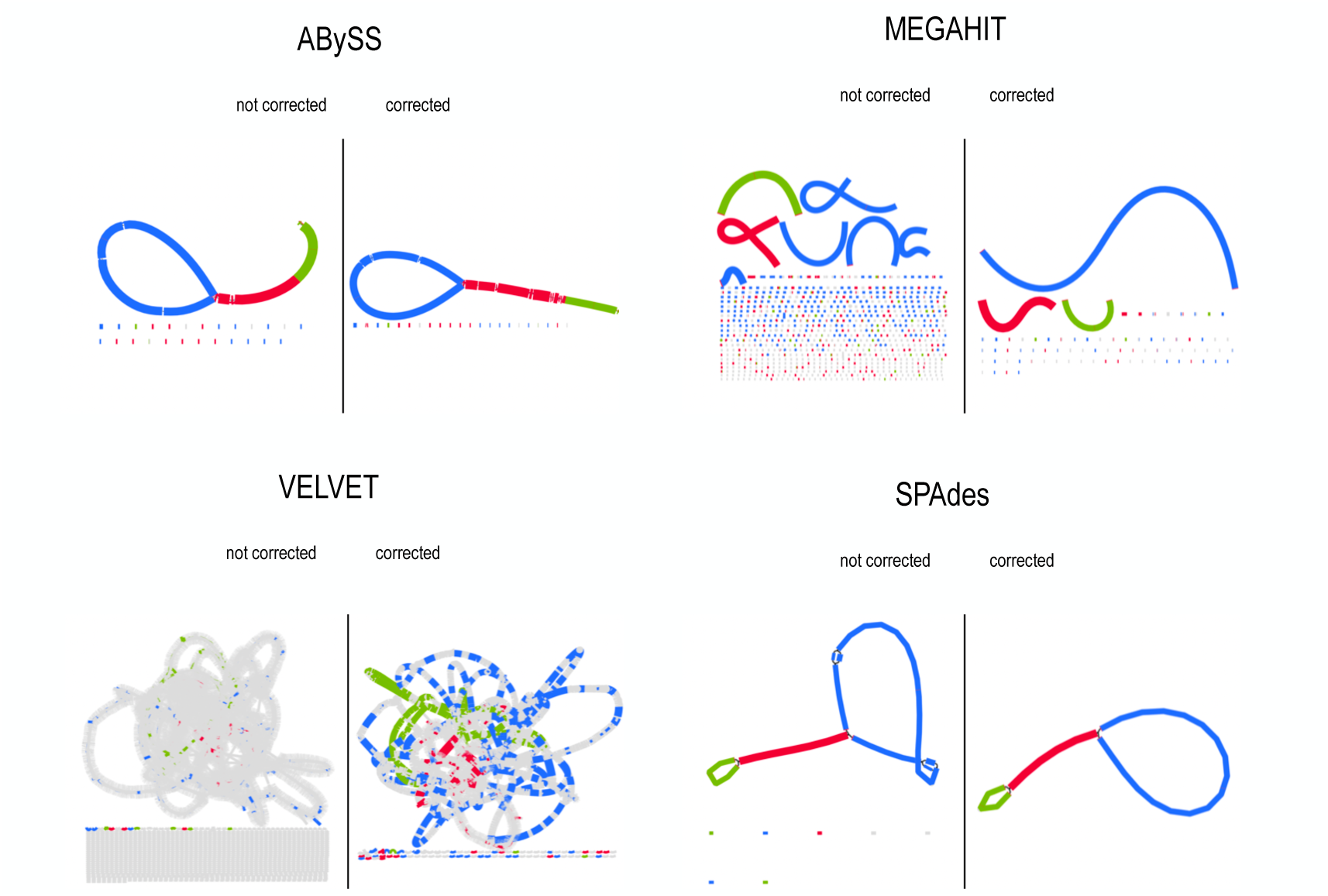
BANDAGE visualization of the L. grandiflora plastome assembly graphs on corrected or uncorrected SRs. Contigs are colored according to their BLAST match to the LSC (blue), SSC (green), and IR (red) segments

**Add. Figure 3:**
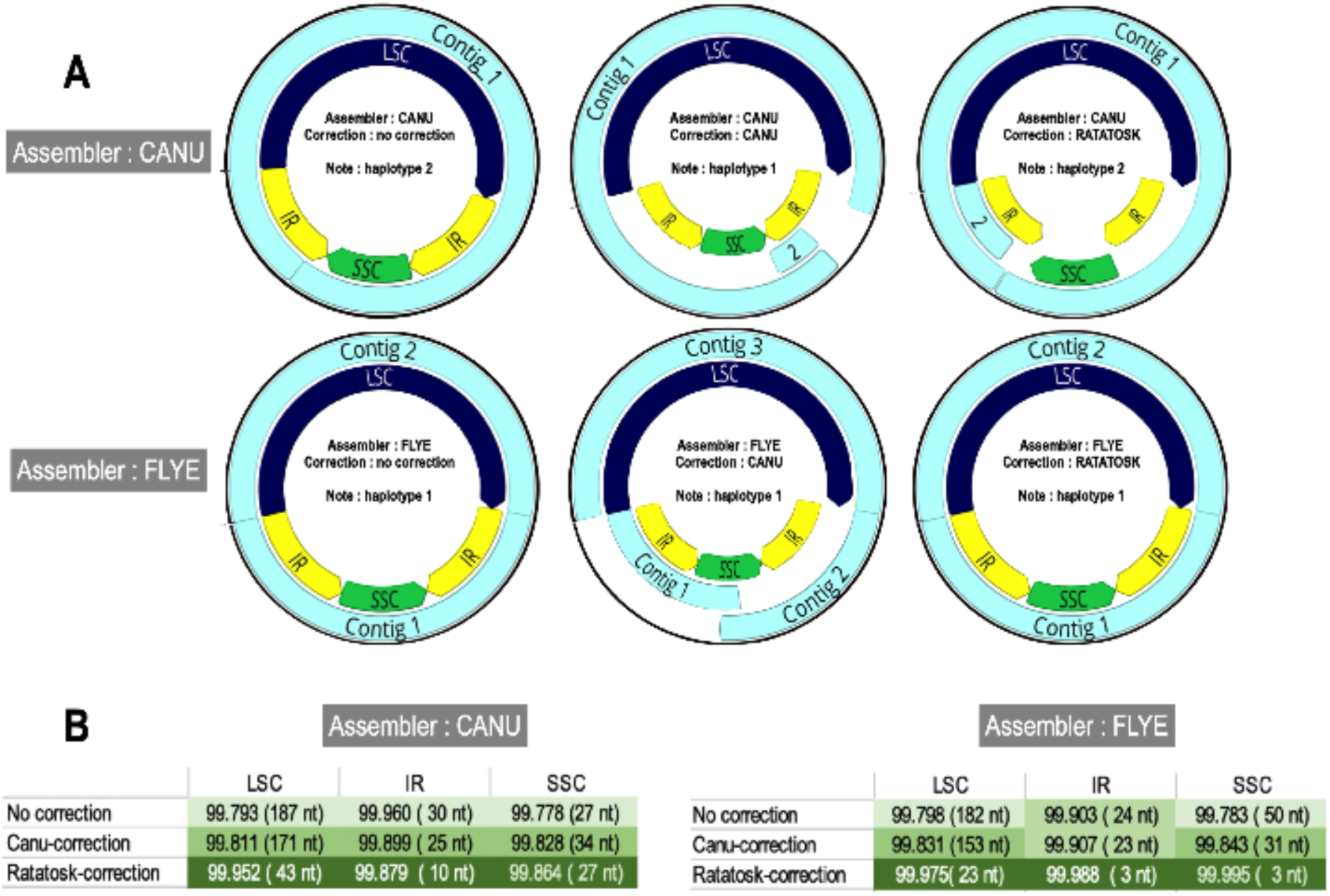
Graphs representing the assemblies of *L. grandiflora* long reads. **A:** Contigs are represented in light blue and the three segments (LSC, SSC and IR) in dark blue, green and yellow, respectively. B: Comparative effectiveness of CANU and RATATOSK correctors.

**Add. Figure 4:**
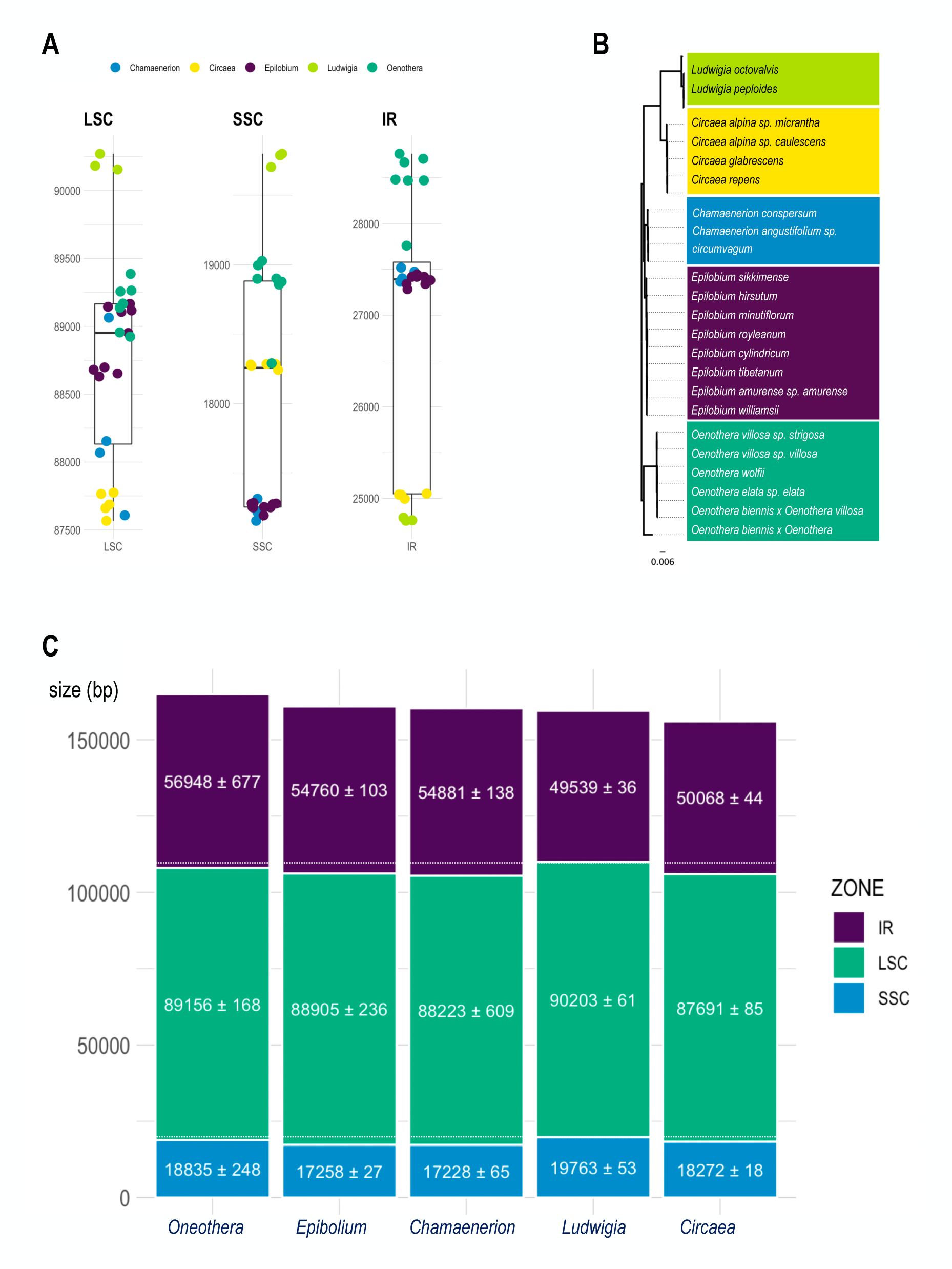
Comparison of LSC, SSC and IR sizes in the Onagraceae. **A:** Comparison of the sizes of LSC, SSC and IR segments in the Onograceae family (*Chamaenerion* in blue, *Circaea* in yellow, *Epibolium* in dark purple, *Ludwigia* in light green and *Oenothera* in dark green). **B:** Maximum likelihood tree made using RAxML (model GTR-GAMMA, algorithm Rapid Hill-climbing) on multiple sequences alignment of Onograceae plastomes made using MAFFT. **C:** Average size of the different chloroplast segments (LSC, SSC and IR) for the 5 genres of Onograceae. IR size corresponds to the sum of the two copies.

**Add. Figure 5:**
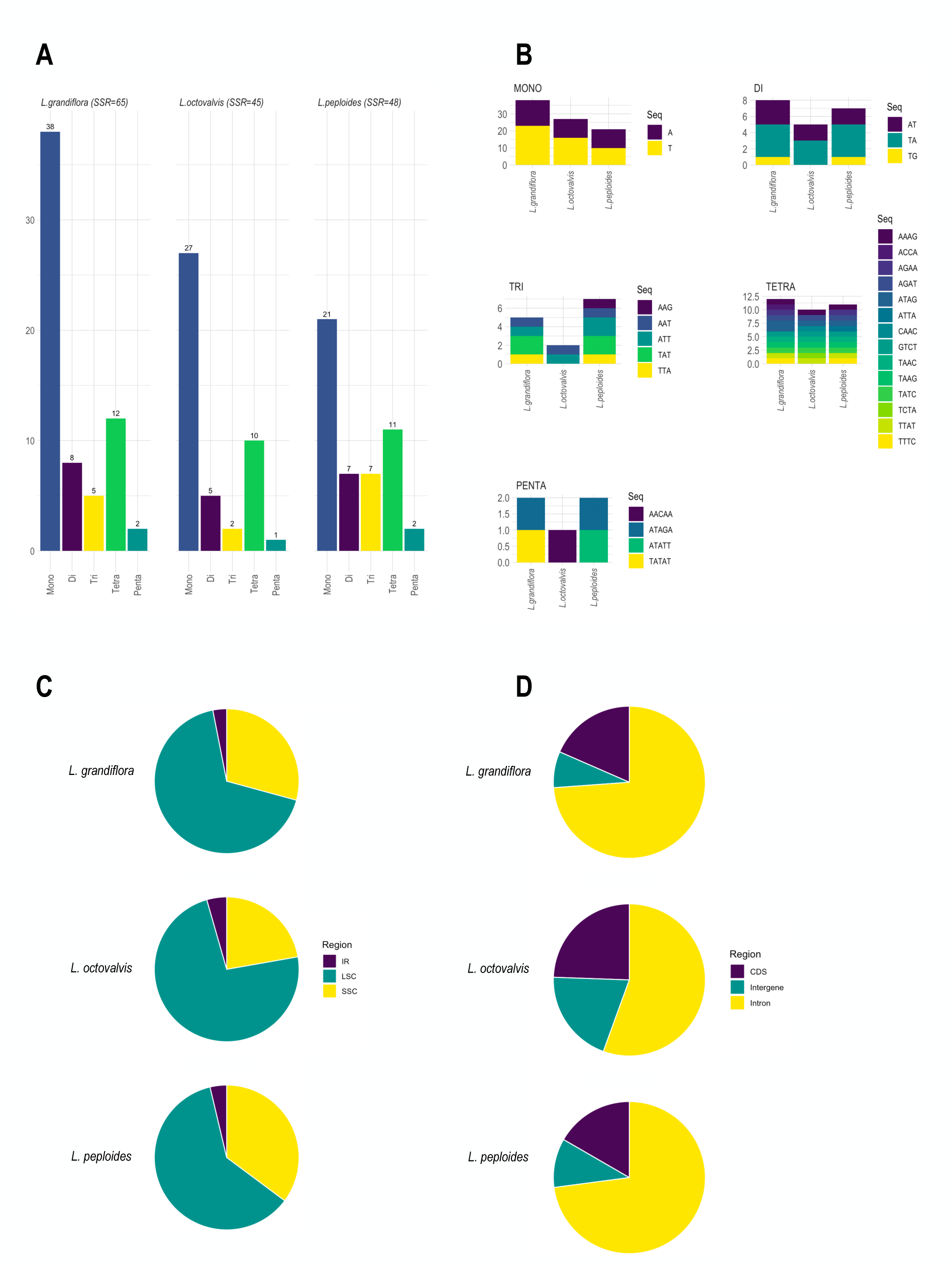
Comparative analysis of Simple-Sequence Repeats (SSRs) in *Ludwigia* chloroplast genomes. A: SSR numbers detected in the three species, by repeat class types (mono, di-, tri-, tetra and pentanucleotides). **B:** Frequency of SSR motifs by repeat class types. **C:** Frequency of SSRs in LSC, SSC and IR regions. **D:** Repartition of SSRs in intergenic, protein-coding and intronic regions.

**Add. Figure 6:**
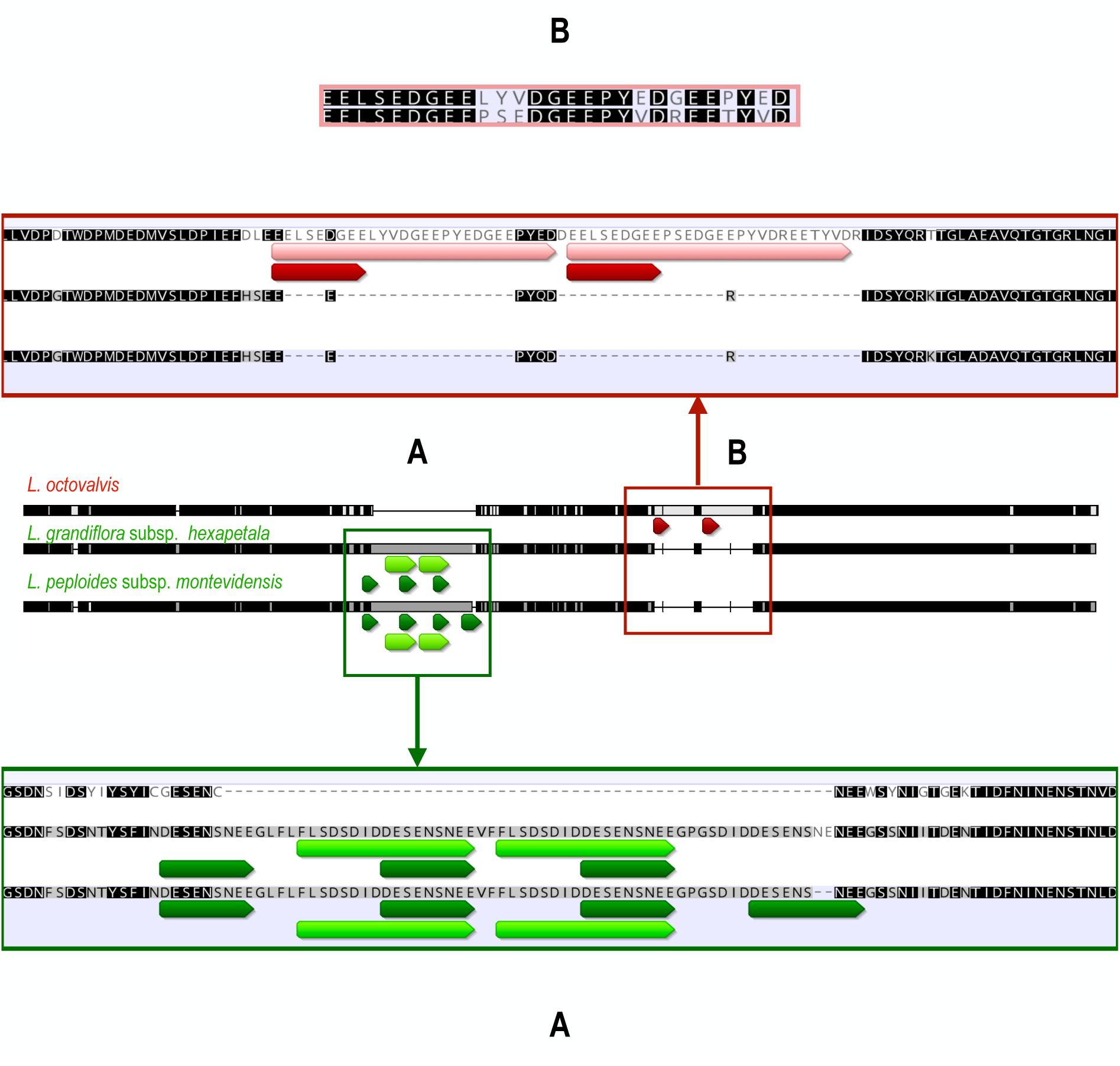
Diagram showing the position of tandem repeats in the accD gene. L. octovalis (in red) and L. peploides and L. grandiflora (in green). We also observe the consequences of these repetitions on the insertion of amino acids, also repeated.

**Add. Figure 7:**
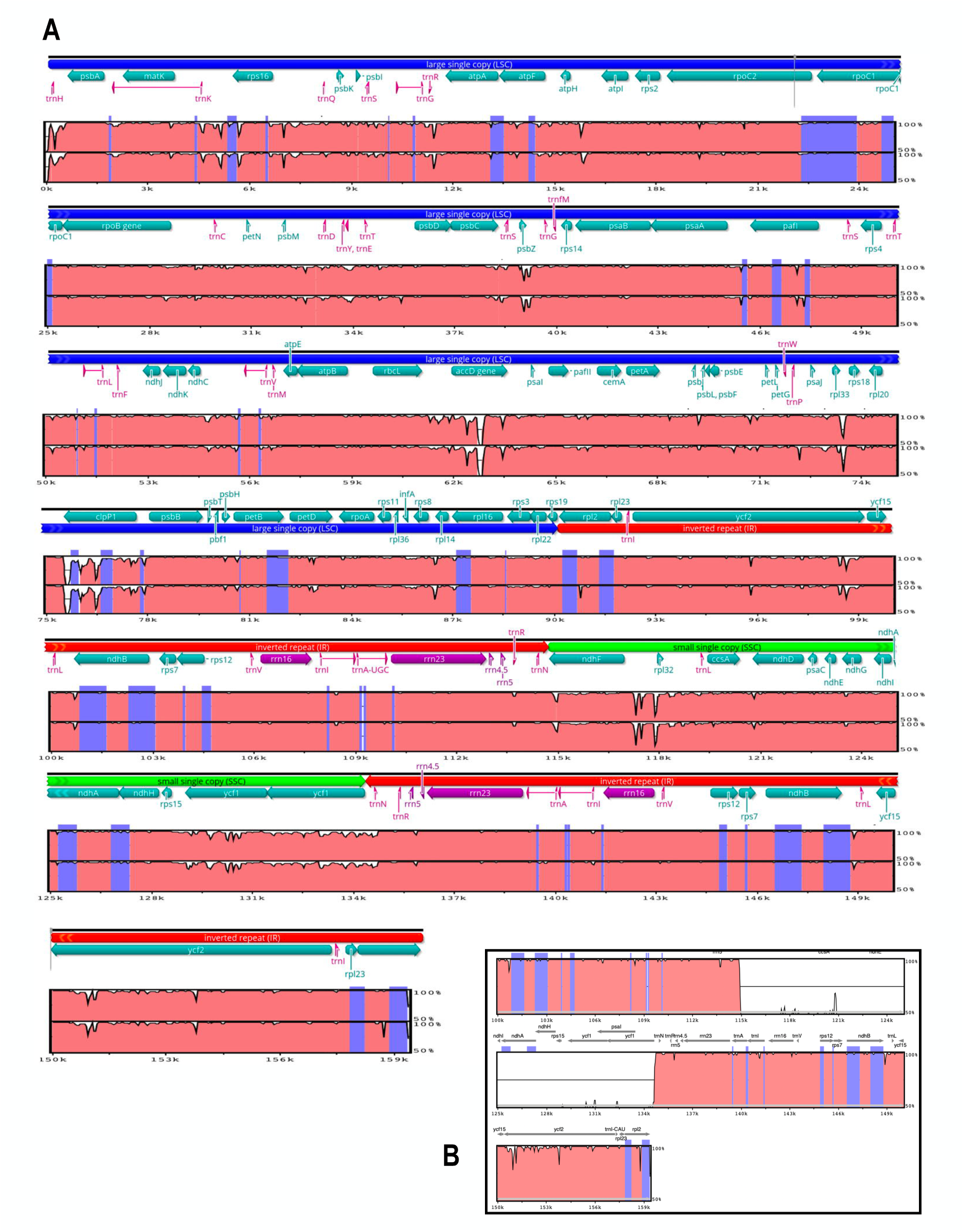
Comparison of the three *Ludwigia* plastomes using mVISTA, with the *L. octovalvis* as a reference. **A:** The y-axis represents the identity percentage (between 50 and 100%). The arrows show the genes (in green: proteins genes, in purple: rRNAs and in fuchsia: tRNAs). Blue blocks indicate exonic regions. LCS, IR and SSC regions are also distinguished (in dark blue, red and green, respectively). The second line corresponds to *L. grandiflora* haplotype 2 (For this haplotype, SSC segment is oriented like *L. octovalvis*) and the third line corresponds to *L. peploides* for which the SSC region has been artificially oriented in the same way as the two other plastomes to allow comparison. **B:** Small box showing a part of the alignment and presenting the consequences if we do not artificially orient the SSC segments in the same direction for the analysis.

**Add. Figure 8:**
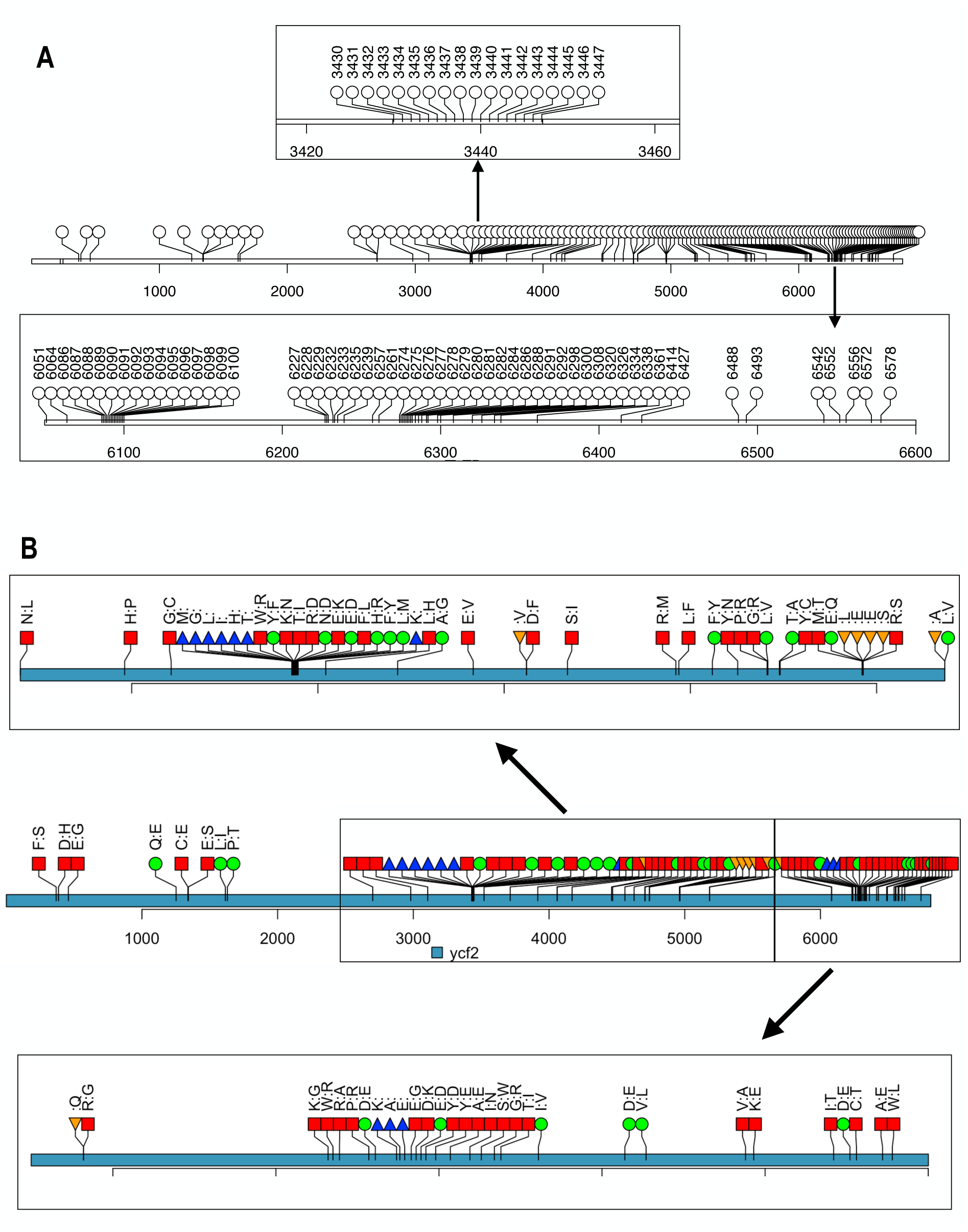
Lollipop diagram allowing the visualization of SNPs and their translational effects on the *ycf2*. **A:** localization of the 256 single nucleotide polymorphisms (SNP) observed by comparing *L. grandiflora*-*L. peploides* with *L. octovalvis.* Two regions particularly dense in SNPs (between 3420 and 3460 and between 6100 and 6600) have been zoomed into to allow better reading. **B:** Effect of these SNPs on the translated sequence of *L. octovalvis*, compared to Ycf2 of the other two species: non conservative mutation: red square; conservative mutation: circle green; deletion: triangle_point_up blue and insertion: triangle_point_down, orange. As for A, two regions were zoomed into in order to distinguish each mutation.

## Notes

### Competing Interest Statement

The authors have declared no competing interest.

### Summary of Updates

we changed the last version of our document because most of the figures and tables were missing.

